# PARP14-dependent ADP-ribosylation of cytoplasmic SQSTM1/p62 foci supports innate immune signaling

**DOI:** 10.1101/2024.06.29.601315

**Authors:** David Kubon, Sudharshana Sundaresan, Diyaraj Nadarajan, Deena M. Leslie Pedrioli, Michael O. Hottiger

**Author notes:** Corresponding author: Michael O. Hottiger.

## Abstract

Interferon-induced ADP-ribosyltransferases are emerging regulators of cellular innate immunity, yet the relevant substrates and downstream effects remain incompletely defined. Here, we mapped the PARP14-dependent ADP-ribosylome in lung epithelial cancer cells and identified the selective autophagy adaptor SQSTM1/p62, along with multiple innate immune regulators, as IFN-dependent MARylation substrates. Site-resolved analysis revealed that PARP14-dependent MARylation is strongly enriched on cysteine residues, including defined cysteine acceptor sites within p62. PARP14-dependent MARylation is localized to cytoplasmic p62 foci that colocalize with diverse ubiquitin species. Furthermore, our results indicate that TRIM21 limits the autophagic degradation of p62, suggesting that the identified p62 foci may have degradation-independent roles. Consistent with this, both PARP14 and p62 support inflammatory gene expression, linking PARP14-dependent MARylation to innate immune regulation. Together, our findings identify p62 foci as sites of PARP14-dependent ADP-ribosylation and suggest a functional link between interferon signaling, ubiquitin-associated protein assemblies, and MARylation-dependent innate immune regulation.

## INTRODUCTION

ADP-ribosyltransferases (ARTs) play an important role in regulating cellular immune responses^1–3^. The diphtheria toxin-like ART subfamily (ARTD) comprises 17 members that catalyze the transfer of ADP-ribose (ADPr) from nicotinamide adenine dinucleotide (NAD^+^) to protein and nucleic acid substrates, resulting in substrate ADP-ribosylation (ADPR)^4,5^. Remarkably, mono-ADP-ribosylating (MARylating) ARTs such as PARP9, 10, 12, 13, and 14 are strongly induced during innate immune responses as a result of the activation of pattern recognition receptors (PRRs) following viral and microbial infection^1,2,6^. However, the relevant substrates and downstream consequences of interferon (IFN)-induced ADPR remain incompletely defined.

Among the group of IFN-inducible mono-ARTs, PARP14 has emerged as a regulator of innate immunity, transcription, DNA replication, and stress responses^7–13^. Recent studies have shown that PARP14 is a dual-function enzyme, exhibiting ADP-ribosyltransferase and -hydrolase activities^14,15^. In particular, the macrodomain 1 of PARP14 functions as an active ADP-ribosylhydrolase^15^. The role of PARP14 in innate immunity has been increasingly recognized, with its identification as one of 38 genes that comprise the interferon-stimulated gene (ISG) signature across tumors^16^. Furthermore, PARP14 activity is regulated by the PARP9/DTX3L complex, positioning PARP14 within a broader IFN-induced ADPR network^17,18^. PARP14 is also one of the few ARTs under positive selection in humans, suggesting an ongoing host-virus conflict^19,20^. Despite growing evidence for a role of PARP14 in host immunity, the substrate specificity and reversal mechanisms of PARP14-dependent ADPR remain poorly understood.

Recent studies reported an increase in punctate cytosolic ADPR foci in human A549 cells upon stimulation with interferons and the dsRNA mimetic poly(I:C)^17,18,21,22^. These foci did not colocalize with known cytoplasmic structures but were strongly enriched for ubiquitin and dependent on PARP14 activity. However, the substrate composition, substrate specificity, structural organization, and functional relevance of these ADPr-positive foci remain only partially resolved.

Selective autophagy receptors are known to aggregate ubiquitin into foci-like structures and can act as organizing platforms that regulate the degradation of ubiquitinated cargo and signaling pathway activity^23,24^. Sequestosome 1 (SQSTM1)/p62 (hereafter p62) is a multifunctional adapter protein involved in ubiquitin-dependent protein sequestration for autophagic clearance and also acts as a signaling platform to control oxidative stress, nutrient sensing, and inflammatory responses^25–28^. p62 contains several domains, including a Phox and Bem1p (PB1) domain, a microtubule-associated protein 1 light chain 3 (LC3)-interacting region (LIR), and a ubiquitin-associated (UBA) domain^29,30^. These domains enable p62 oligomerization, ubiquitin binding, and recruitment to the autophagy machinery, promoting the formation of droplet-like cytoplasmic p62 foci^31,32^. The composition and function of these foci are shaped by multiple posttranslational modifications and binding partners, which tune p62 behavior toward distinct degradation-dependent and signaling-related functions^27,31^.

Here, we identify p62 as a type I IFN-induced, PARP14-dependent ADPR target in lung epithelial cells. Evidence shows that PARP14-dependent ADPR predominantly targets cysteine residues, including specific cysteine sites on p62. This ADPR localizes to cytoplasmic p62 foci, which are distinct from canonical autophagosomes. Our findings suggest these foci serve as PARP14-dependent ADPR hubs, with autophagic turnover of p62 being inhibited by TRIM21. Collectively, these results support a model in which PARP14-dependent ADP-ribosylation marks p62 foci that are maintained for degradation-independent functions and may contribute to innate immune regulation.

## RESULTS

### SQSTM1/p62 is a major ADP-ribosylated target under inflammatory conditions

To identify mono-ART-driven ADPR in response to IFN-induced innate immune signaling, we used A549 lung epithelial cancer cells. This model exhibits low basal expression of interferon-stimulated genes (ISGs) and low levels of ADPR but shows robust induction of ISGs and ADPR upon treatment with interferons and the dsRNA mimetic poly(I:C)^22^. A549 cells were thus treated for 24 hours with IFN-β, a type I interferon that results in robust induction of IFN-responsive mono-ARTs^2^. We first analyzed the IFN-β-induced ADP-ribosylome using an established ADP-ribose-specific mass spectrometry (MS)-based proteomics workflow^33,34^. We observed a strong change in ADP-ribosylated proteins (i.e., the ADP-ribosylome) in A549 cells comparing basal (untreated) to IFN-β-treated conditions. The modification profile shifted from predominantly detecting PARP1 and histones towards modifications on PARP12, DTX3L, and p62 after IFN-β treatment (Fig. S1A). While some physiological ADPr sites on PARP12 have been previously described^34^, both DTX3L and p62 emerged as novel proteins modified by ADPr. Initial site-mapping identified p62 ADPR on cysteine residues C113 and either C289 or C290 (Fig. 1B) in A549 cells. In addition, a modification at C331 was identified in NCI-H1975^35^. Because recent advances in ADP-ribosylation proteomics have shown that conventional workflows can underestimate chemically labile ADPR acceptor sites, particularly on acidic residues such as aspartate and glutamate^36,37^, we performed a second IFN-β-induced ADP-ribosylome using an optimized workflow to preserve labile ADPR sites. This approach substantially increased the number of detected ADP-ribosylated proteins and sites (Fig. 1A). Importantly, p62 again emerged as a major IFN-induced ADPR target. The expanded dataset confirmed cysteine ADPR on p62 and identified additional acceptor sites, including C128, C331, D108, S294, and R110. Thus, p62 is modified at different acceptor sites, with cysteine ADPR being a prominent and reproducible component of its IFN-induced modification pattern. The identified cysteine sites are positioned near functionally relevant regions of p62, including the ZZ-type zinc finger domain, the nuclear localization and export sequences, and the LC3-interacting region. Given the established roles of p62 in selective autophagy, inflammatory signaling, antiviral defense and the reproducible detection of IFN-induced p62 ADPR at these sites, made p62 a strong candidate for mechanistic follow-up.

**Figure 1:**
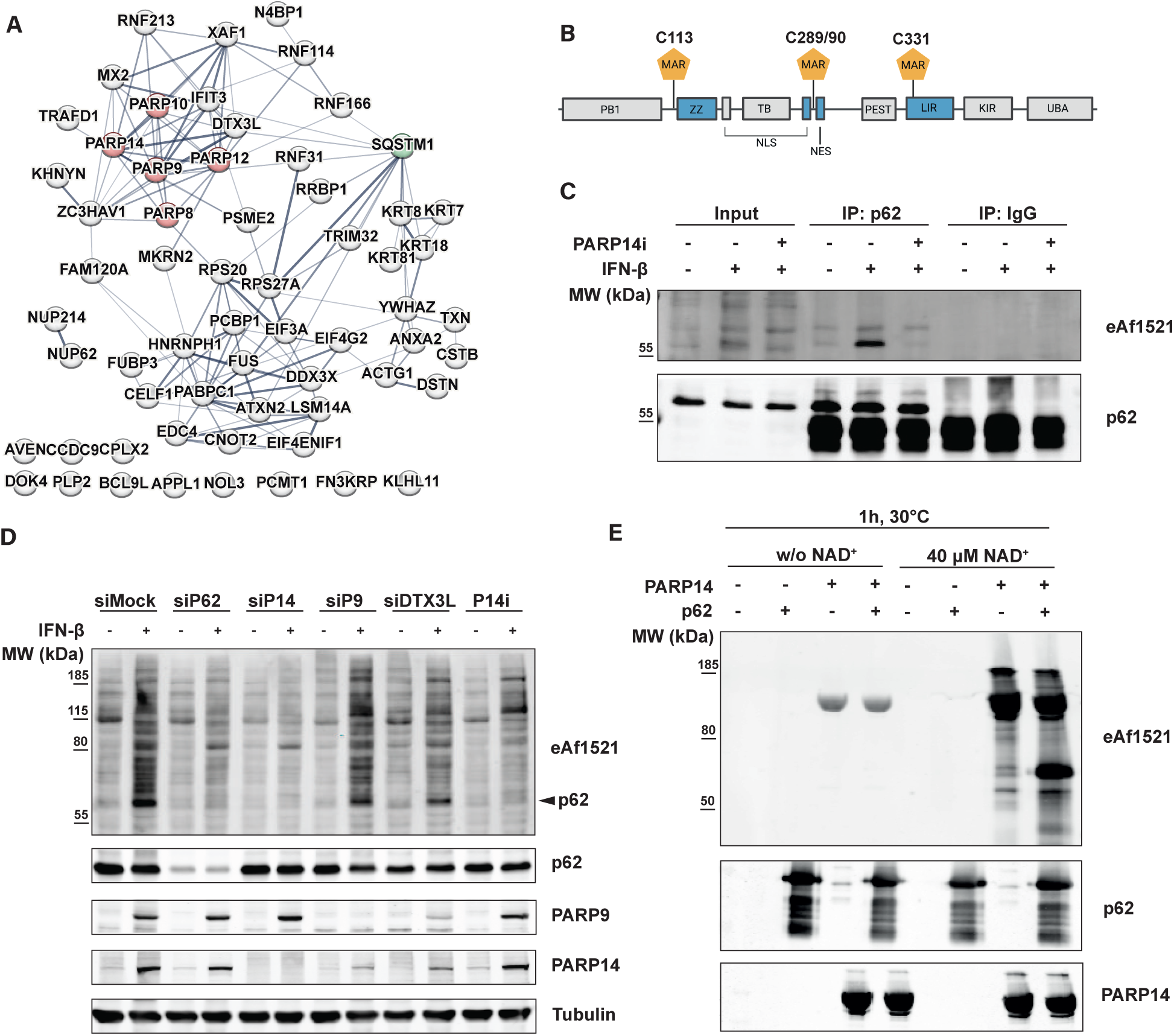
SQSTM1/p62 is ADP-ribosylated at cysteine residues in a PARP14-dependent manner. **(A)** STRING analysis of ADP-ribosylated proteins identified by LC-MS/MS in A549 cells treated for 24 h with IFN-β (1000 U/mL), in comparison to untreated control cells. ADP-ribosyltransferases are marked in red, SQSTM1/p62 is marked in green. **(B)** p62 domain structure with annotation of the dominant cysteine ADPR sites identified by LC-MS/MS. Figure was created with a licensed version of BioRender.com (License number for Fig. 1B: LV26ZULJXG) **(C)** p62 immunoprecipitation from A549 cells treated with IFN-β (1000 U/mL) and the PARP14 inhibitor (P14i) RBN012759 (20 nM) for 24 h. **(D)** Immunoblots of A549 cells following the knockdown of p62, PARP14 (siP14), PARP9 (siP9), and DTX3L for 48 h. IFN-β and PARP14i treatment according to Fig. 1C. **(E)** Immunoblot of *in vitro* reaction using recombinant full-length PARP14 and recombinant full-length p62. Reaction was performed in the presence of 40 μM NAD^+^ for 1 h at 30°C.

### Cysteine residues are major IFN-induced ADP-ribosylation sites on p62

To corroborate the MS results, we immunoprecipitated endogenous p62 and detected its ADPR using the engineered macrodomain eAf1521. Basal levels of p62 ADPR were minimal, whereas IFN-β treatment significantly increased the amount of modified p62 (Fig. 1C). The IFN-β-dependent induction of p62 ADPR after treatment suggests that the modification is catalyzed by one of the IFN-inducible mono-ARTs, which are strongly upregulated upon IFN-β treatment (Fig. S1B). This possibility is further supported by the observation that p62 ADPR levels do not increase immediately but start to appear after 8 hours (Fig. S1C). Notably, this modification was induced not only by IFN-β but also by the type II interferon IFNγ and poly(I:C) (Fig. S1D), indicating broad functional consequences given its induction by multiple stimuli. To functionally validate the major MS-identified acceptor sites, we focused on the cysteine residues initially detected in p62. We mutated the corresponding cysteine residues and complemented A549 cells with a doxycycline-inducible siRNA-resistant HA-tagged ADPR-deficient p62 mutant (ΔADPR). Expression of the p62 mutant, while simultaneously depleting endogenous p62, revealed a loss of the p62 ADPR signal upon overexpression of the ADPR-deficient mutant compared to wildtype p62 (Fig. S1F). These data identify p62 as a major IFN-induced ADPR target and establish the mapped cysteine residues as major ADPR modification sites on p62.

### PARP14 activity is required for IFN-induced SQSTM1/p62 ADP-ribosylation

Several mono-ARTs are induced by IFN signaling^2^ and could contribute to p62 ADPR. We focused on PARP14 because the timing and magnitude of PARP14 up-regulation closely coincided with the observed p62 ADPR signal (Fig. S1C and S1E). Additionally, PARP14 has emerged as a major driver of IFN-induced ADPR^11,13,38^. Immunoprecipitation of endogenous p62 followed by ADPR detection using immunoblotting and eAf1521^33^, revealed a complete loss of p62 modification upon combinatorial treatment with IFN-β and the selective PARP14 inhibitor RBN012759 (i.e., PARP14i)^9^ (Fig. 1C). Consistent with pharmacological inhibition, siRNA-mediated depletion of PARP14 also abolished IFN- β-induced p62 ADPR (Fig. 1D). These results indicate that PARP14 is directly or indirectly (e.g., through another ART) responsible for the modification of p62. To investigate these possibilities, we performed a knockdown of PARP9 and its binding partner, DTX3L, which have recently been described as forming a complex with PARP14 and cross-regulating each other’s protein levels^18,38^. Neither knockdown of PARP9 nor DTX3L affected the ADPR levels of p62, whereas siRNA-mediated depletion or inhibition of PARP14 abolished p62 ADPR (Fig. 1D and S1G). To test whether PARP14 is sufficient to modify p62, we performed an *in vitro* ADPR assay using recombinant full-length PARP14 and recombinant full-length p62 in the presence of NAD⁺. Under these conditions, PARP14 showed strong auto-ADPR and promoted trans-ADPR of p62 (Fig. 1E). Together, these results identify PARP14 as the major upstream enzyme responsible for IFN-β-induced p62 ADPR and suggest that p62 is a direct PARP14 substrate.

Given the role of PARP14 in antiviral immunity and the ability of viral macrodomains to antagonize host ADPR^39–41^, we tested whether p62 ADPR is sensitive to SARS-CoV-2 nsP3 Mac1 activity. Overexpression of SARS-CoV-2 nsP3 Mac1 reduced IFN-β-induced p62 ADPR in cells (Fig. S1H and S1I), and recombinant Mac1 suppressed both p62 and PARP14 modification in a reconstituted PARP14-p62 *in vitro* assay (Fig. S1J). These results suggest that PARP14-dependent p62 ADPR is sensitive to Mac1 activity. However, the experiments do not distinguish between direct hydrolysis of ADPr from p62 and indirect suppression of p62 trans-modification through PARP14 de-modification.

### PARP14-dependent ADP-ribosylation localizes to cytoplasmic SQSTM1/p62 foci

Given that p62 emerged as both a major PARP14-dependent ADPR target and a potential scaffold due to its oligomerization capabilities^32,42^, we investigated whether IFN-induced ADPR is spatially associated with p62-positive cytoplasmic structures. A549 cells were therefore treated with IFN-β, both alone and in combination with the PARP14i, followed by immunofluorescence analysis. These experiments revealed that p62 exclusively localized to the cytoplasm, forming two distinct pools: phase- separated foci^32^ and a dispersed form throughout the cytoplasm (Fig. 2A). We observed an increase in ADPR foci upon IFN-β treatment, which colocalized with p62 foci (Fig. 2A and 2B). Notably, neither total p62 levels nor the number of p62 foci showed any discernible change in response to different treatment conditions (Fig. S2A and S2B), indicating that PARP14-dependent ADPR does not affect the formation of p62 foci. These findings suggest that p62 is a major ADP-ribosylated component of the foci, while also leaving open the possibility that additional ADP-ribosylated proteins are recruited to or modified within the same structures. p62 shuttles between the cytoplasm and nucleus, thereby facilitating the degradation of ubiquitinated proteins in both compartments. Given the proximity of the ADPR sites C289/290 to the nuclear localization signal (NLS) and the nuclear export signal (NES)^43^ (Fig. 1B), we investigated whether ADPR affects p62 localization. Cells were thus treated with the nuclear export inhibitor leptomycin B, which traps p62 in the nucleus. After 4 hours of treatment, p62 was found exclusively in the nucleus, unaffected by previous treatment with IFN-β and PARP14i (Fig. 2C). These findings indicate that PARP14-dependent ADPR does not affect p62 shuttling behavior.

**Figure 2:**
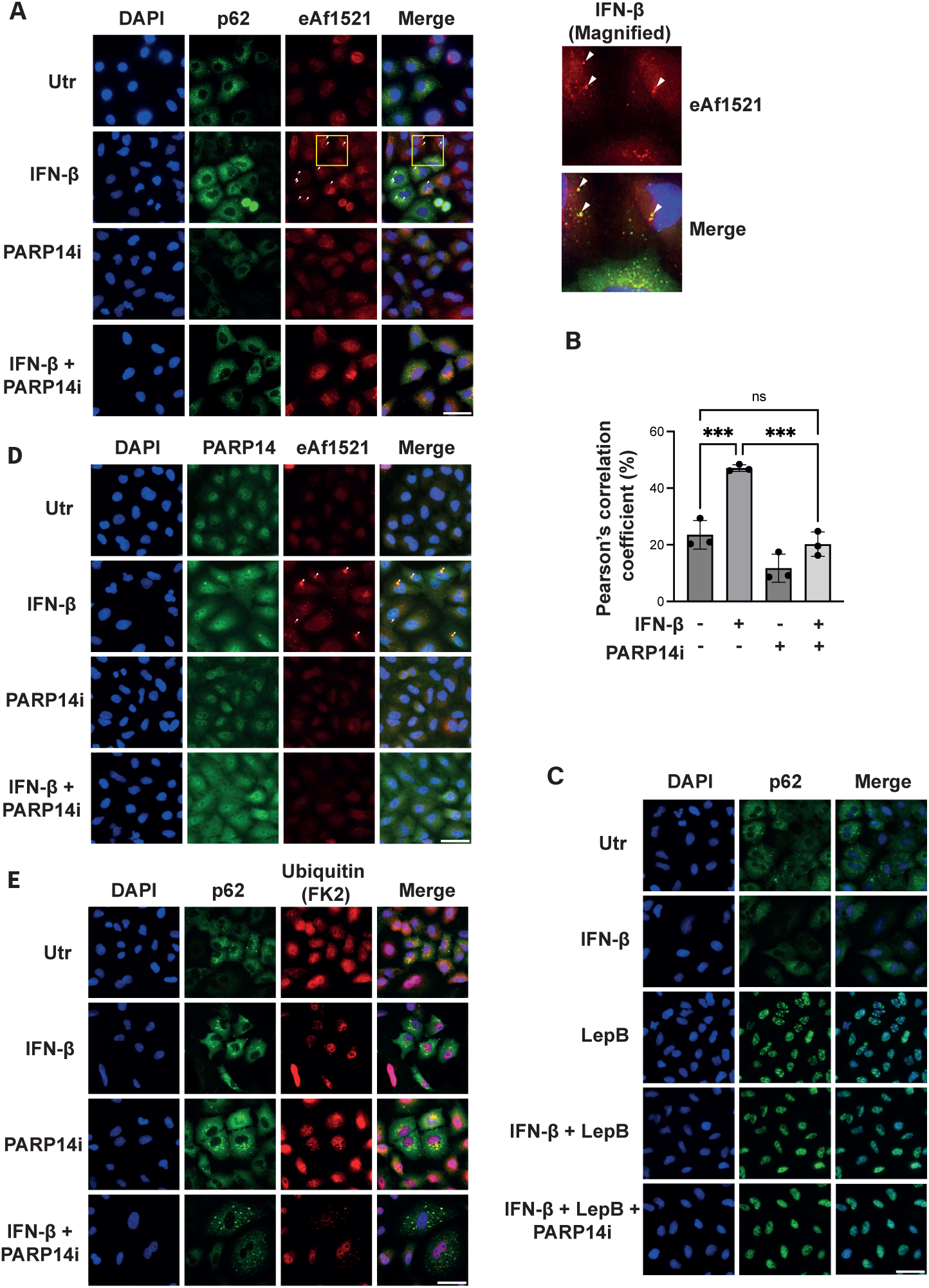
SQSTM1/p62 forms cytoplasmic foci which colocalize with ADP-ribosylation, PARP14 and ubiquitin. **(A/B)** A549 cells were treated with IFN-β and PARP14i (20 nM) for 24 h, and p62 localization and its colocalization with ADPR foci were analyzed by immunofluorescence. Representative images **(A)** and quantification of p62 and ADPR foci colocalization **(B)** are shown. White arrows indicate colocalizing foci. **(C)** Representative immunofluorescence images of A549 cells treated according to Fig. 2A/B. For the last 4 h of the treatment, co-treatment with leptomycin B (LepB) (30 nM) was performed. **(D)** PARP14 and ADPr foci colocalization was analyzed by immunofluorescence in A549 cells treated according to Fig. 2A/B. Representative images are shown. **(E)** p62 and ubiquitin colocalization was analyzed by immunofluorescence in A549 cells treated according to Fig. 2A/B. One-way ANOVA determined P values for experiments with more than three conditions and multiple comparisons analysis using Tukey’s multiple comparison test. * p<0.05, ** p<0.005, *** p<0.0005, ns not significant. The scale bar represents 20 μm.

### PARP14-dependent ADP-ribosylation is enriched on cysteine residues and modifies ubiquitin- and innate immune-regulatory proteins

Although p62 was strongly ADP-ribosylated after IFN-β stimulation, mutations of the major p62 acceptor sites did not completely abolish the ADPR signal within p62 foci (Fig. S2F). This suggested that p62 itself is unlikely to account for the entire ADPR signal within these structures and raised the possibility that PARP14 modifies additional targets. Our initial ADP-ribosylome analysis identified IFN-induced ADPR targets but did not define which modification events were specifically dependent on PARP14 activity. We therefore performed a second ADP-ribosylome experiment designed to map the PARP14-dependent ADPR landscape under inflammatory conditions using pharmacological PARP14 inhibition. We again applied our improved MS workflow, which preserves labile ADPR events, in combination with unbiased eAf1521 enrichment (Fig. 3A-E). A549 cells were treated with IFN-β in the presence or absence of the selective PARP14 inhibitor RBN012759, and ADP-ribosylated peptides were analyzed by mass spectrometry. This analysis confirmed p62 as a major PARP14- dependent target. PARP14 inhibition strongly reduced the main IFN-β-induced p62 modification sites, including C113, C289, C290, C331, D108 and R110. Although multiple acceptor sites were detected, cysteine residues represented the dominant PARP14-dependent acceptor sites on p62 under these conditions. Consistent with this observation, global analysis of the PARP14-dependent ADP- ribosylome showed that PARP14 inhibition preferentially reduced cysteine ADPR. Approximately 83% of the ADPR sites reduced upon PARP14 inhibition were mapped to cysteine residues. In contrast, the next most frequent acceptor class comprised the acidic amino acids glutamate and aspartate, accounting for approximately 8% of reduced sites (Fig. 3D). These data suggest that PARP14-dependent ADPR under IFN-β-stimulated conditions mainly affects cysteine residues.

**Figure 3:**
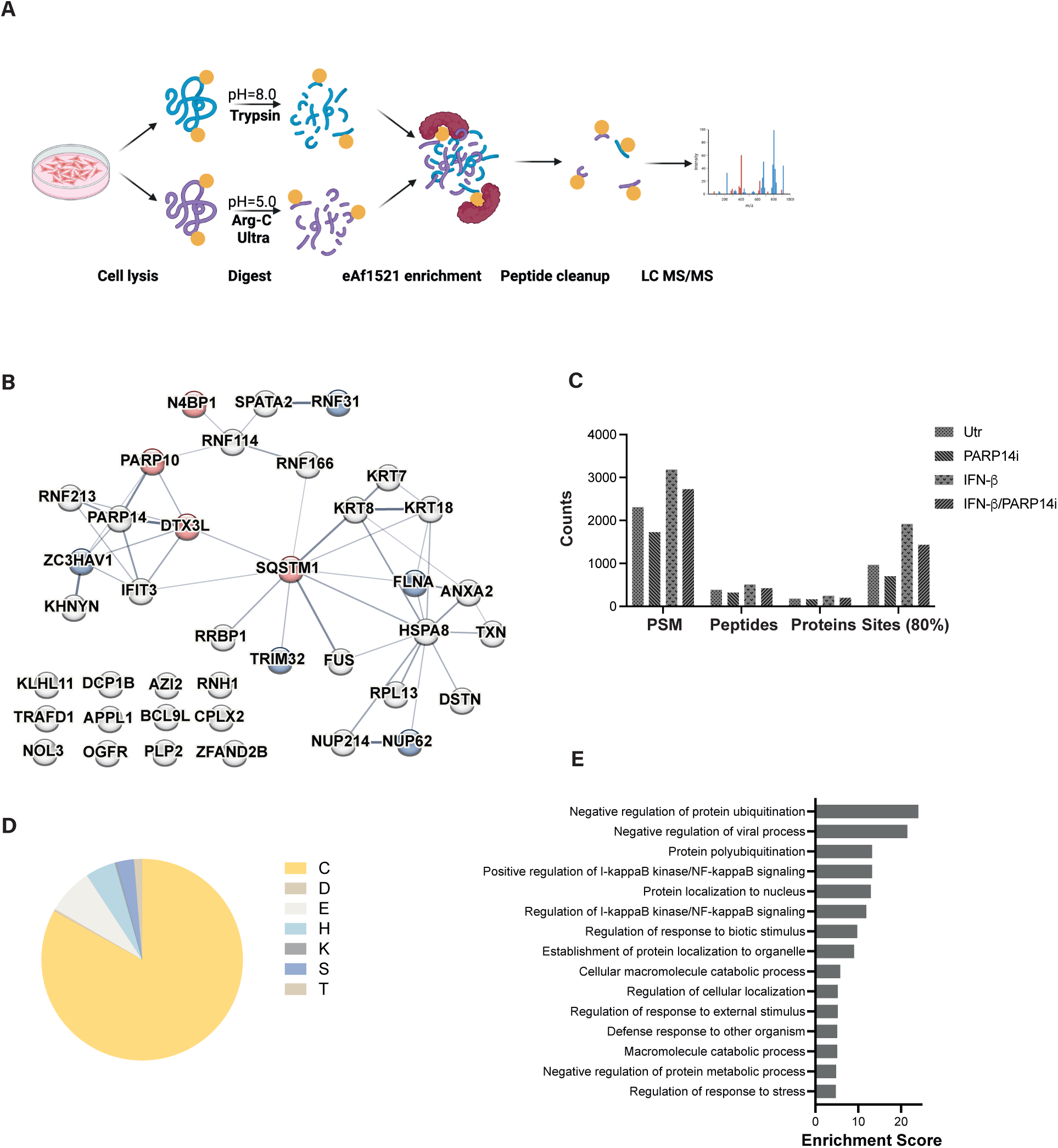
PARP14 modifies innate immune factors and ubiquitin regulatory proteins on cysteine residues. **(A)** Diagram of our optimized ADP-ribosylome workflow with a hybrid tryptic/Arg-C Ultra digest at different pH conditions followed by enrichment of modified peptides with eAf1521 and LC-MS/MS analysis. Figure was created with BioRender.com **(B)** STRING analysis of PARP14-dependent ADPR targets identified by LC-MS/MS using the PARP14i. Nodes marked in red correspond to the GO term ‘Negative regulation of protein ubiquitination’, nodes marked in blue correspond to the GO term ‘Positive regulation of I-kappaB kinase/NF-kappaB signaling’. **(C)** Bar chart showing the summary metrics of the performed ADP-ribosylome across the different measured conditions. Peptide-spectrum matches (PSMs), unique ADPR peptides, proteins, and localized ADPR sites (80% confidence threshold). **(D)** Pie chart showing the PARP14-dependent acceptor site preference identified from our PARP14-dependent ADP-ribosylome. The one-letter amino acid code is displayed. **(E)** Bar chart showing the enrichment score (FDR < 0.05) of the GO analysis (Biological process) based on the PARP14-dependent ADPR targets identified.

Functional enrichment analysis of the PARP14-dependent ADP-ribosylome revealed prominent enrichment for terms related to antiviral defense, ubiquitination regulation, and NF-κB signaling (Fig. 3E). Thus, rather than representing an isolated p62 modification event, PARP14-dependent ADPR appeared to affect a broader network of proteins linked to innate immunity and ubiquitin-dependent protein regulation, providing potential insight into the pathways associated with p62-positive ADPr foci. Within this network, two candidate clusters were particularly notable for their role in connecting PARP14-dependent ADPR to established innate immune effector pathways. The first contained the antiviral RNA restriction factors PARP13/ZAP, KHNYN, and N4BP1^44–46^, which have been shown to cooperate to degrade viral RNA^44,47^. A second cluster includes RNF31, also known as HOIP, the catalytic E3 ligase component of the linear ubiquitin chain assembly complex (LUBAC)^48^. RNF31 mediates M1-linked linear ubiquitination and regulates NEMO-dependent NF-κB signaling^49^. In addition, SPATA2 was identified as a PARP14-dependent ADPR target. SPATA2 bridges the deubiquitinase CYLD to LUBAC, thereby modulating M1-linked ubiquitin signaling^50^. The dataset also contained RNF114 and RNF166, two E3 ubiquitin ligases recently described as mediators of ubiquitin- ADP-ribose hybrid mark extension through K11-linked ubiquitination and proteasome-dependent degradation^51–54^. Together, these data suggest that PARP14-dependent ADPR converges on several interconnected proteostasis and innate immune pathways, including ubiquitin signaling, protein degradation, antiviral RNA restriction, and regulation of the NF-κB pathway. In this broader inflammatory ADPR program, p62 emerges as a major substrate and potential scaffold for PARP14- dependent activity.

### PARP14 and ubiquitin colocalize with ADP-ribosylated SQSTM1/p62 foci

Having established that the PARP14-dependent ADP-ribosylome is enriched for antiviral, inflammatory, and ubiquitin-regulatory proteins, we further characterized the identity of the detected p62 foci. We first co-stained IFN-β-treated A549 cells for markers of major cytoplasmic compartments. These experiments revealed that neither markers for mitochondria, autophagy, nor the endoplasmic reticulum (ER) were enriched in p62 foci (Fig. S2C-E), indicating that the ADP-ribosylated p62 foci represent a distinct p62 pool within the cell.

Because the PARP14-dependent ADP-ribosylome was enriched for ubiquitin pathway components and p62 functions as a scaffold for ubiquitinated cargo, we assessed whether these foci associate with ubiquitin. Immunofluorescence analysis revealed that the p62 foci are ubiquitin-positive (Fig. 2E) but independent of their ADPR status, suggesting that the ubiquitination is not induced by the ADPR of p62 by PARP14. Similarly, M1-linked ubiquitination was identified in p62-positive foci (Fig. S2G), but this signal was not affected by PARP14 inhibition. We next investigated whether PARP14 itself colocalizes with the observed p62 foci, by treating A549 cells with IFN-β alone and in combination with the PARP14i (Fig. 2D). The modified foci colocalized with PARP14 in an IFN-β-dependent manner, supporting its role as the primary writer of p62 ADPR. Interestingly, treatment of cells with PARP14i dissociated PARP14 from the foci, suggesting that the enzymatic activity of PARP14 is required for its recruitment or retention in p62 foci (Fig. 2D). Together, our data indicate that p62 foci form independently of PARP14-dependent ADPR.

### ADP-ribosylated SQSTM1/p62 remains autophagy-competent but is protected from turnover by TRIM21

Selective autophagy is considered one of the primary functions of p62. The enrichment of ubiquitin within these structures also pointed toward a degradative role of these foci. Although p62 foci did not colocalize with LC3 under steady-state conditions, autophagy blockage using the lysosomotropic agent chloroquine (CQ), which inhibits the fusion and maturation of endosomes and lysosomes with autophagosomes^55^, resulted in accumulation of both total p62 and its ADP-ribosylated form (Fig. 4A). This observation suggests that, despite the absence of LC3 accumulation at p62 foci, ADP-ribosylated p62 can undergo autophagy-dependent degradation. GFP-Trap pull-down of overexpressed LC3-EGFP confirmed that both p62 and its ADP-ribosylated form interact with LC3 (Fig. 4B). In line with these findings, global proteome analysis comparing IFN-β-treated cells with IFN-β+PARP14i treated cells revealed only minor changes in overall protein abundance (Fig. S3A). We further tested whether p62 ADPR affects selective autophagic targeting of viral proteins. For this, we overexpressed hepatitis C virus core protein, which has previously been reported to undergo p62-dependent autophagic degradation^56^. Consistent with this, depletion of p62 partially stabilized the viral core protein, supporting a contribution of p62 to its turnover (Fig. S3B). However, PARP14 knockdown did not produce a comparable stabilization phenotype. These data suggest that p62 contributes to the degradation of this viral cargo but in a PARP14-independent manner.

**Figure 4:**
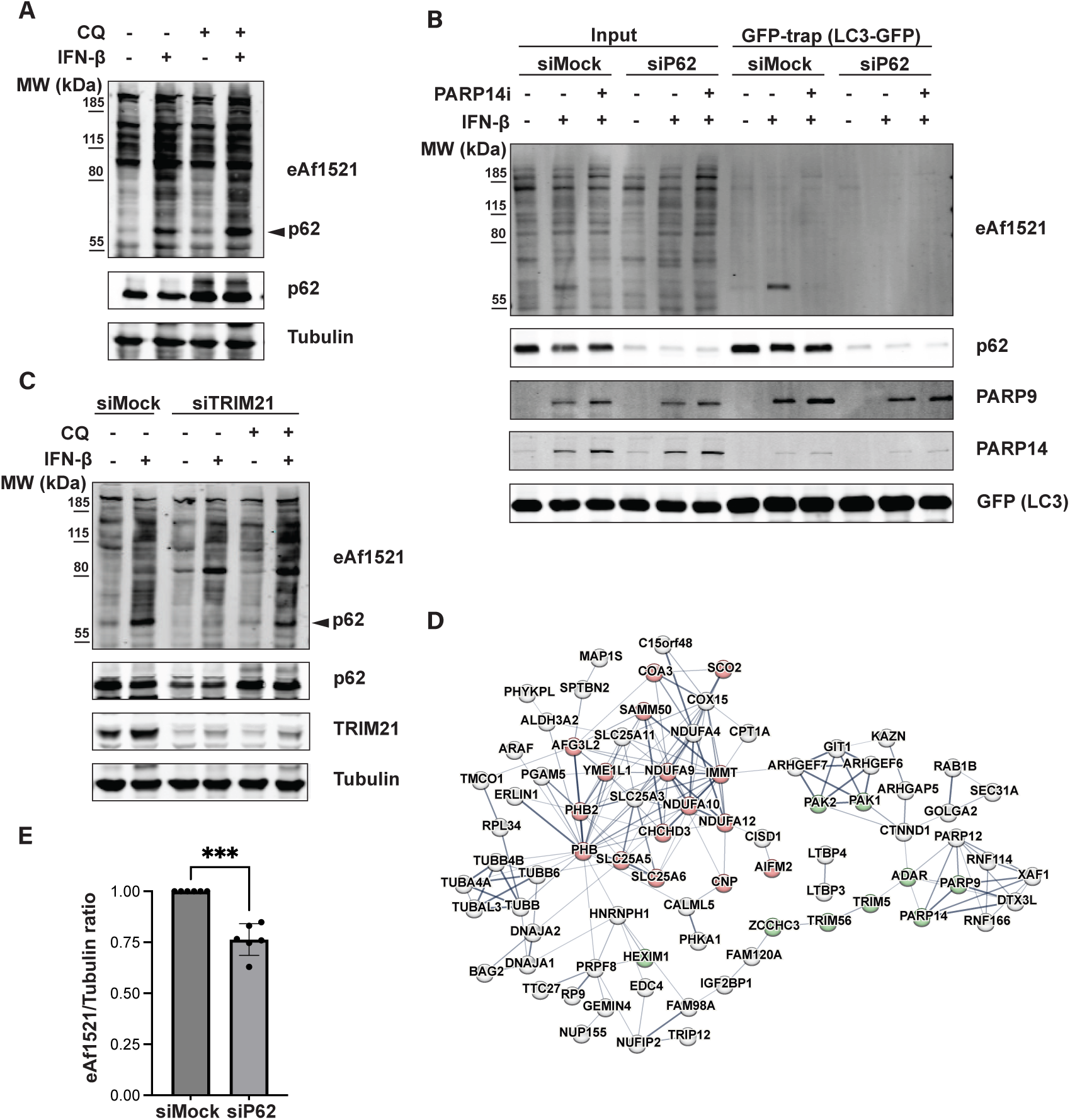
Autophagic turnover of ADP-ribosylated p62 is prevented by TRIM21. **(A)** A549 cells were treated with IFN-β (1000 U/mL) and 10 μM of chloroquine (CQ) for 24 h, followed by immunoblot analysis. **(B)** GFP-Trap co-immunoprecipitation of A549 cells with stable overexpression of a doxycycline-inducible LC3-EGFP construct and siRNA knockdown of p62 (48 h). For the last 24 h of the experiment, cells were treated with 500ng/mL doxycycline to induce LC3-EGFP expression, IFN-β (1000 U/mL), and PARP14i (20 nM). **(C)** Immunoblot of A549 cells after TRIM21 knockdown for 48 h. Cells were co-treated with IFN-β (1000 U/mL) and chloroquine (CQ) (10 μM) for the last 24 h of the experiment. **(D)** STRING analysis of co-IP MS experiment showing proteins more strongly enriched following IFN-β+PARP14i treatment in comparison to IFN-β alone. Endogenous p62 was immunoprecipitated from A549 cells treated with IFN-β (1000 U/mL) and PARP14i (20 nM) for 24 h. Nodes marked in red correspond to the GO term ‘Mitochondrion organization’, green nodes correspond to the GO term ‘Regulation of innate immune response’. **(E)** Quantification of immunoblots from Fig. 1D and S1D comparing the whole lane eAf1521 signal between IFN-β-treated conditions (siMOCK) and IFN-β + siP62. P-values were determined by a paired, two-tailed Student’s t-test.

Contrary to our expectations, we observed no significant changes in p62 protein levels upon IFN- β treatment (Fig. 4A), although the existing literature suggests that type I interferons enhance autophagy to increase viral clearance and antigen presentation^57,58^. This suggests that IFN-induced mechanisms may prevent p62 from undergoing autophagic clearance. The tripartite motif-containing protein 21 (TRIM21), which is induced by IFN treatment in A549 cells, has been shown to mediate p62 ubiquitination via its PB1 domain, thereby preventing self-oligomerization and autophagic targeting^59,60^. We therefore depleted TRIM21 to test whether it regulates the stability of ADP-ribosylated p62. TRIM21 knockdown reduced both the total and ADP-ribosylated p62 (Fig. 4C). This reduction was rescued by co-treatment with CQ, indicating that loss of TRIM21 promotes autophagic degradation of p62, including its ADP-ribosylated form. TRIM21 does not appear to selectively protect only the ADP-ribosylated form of p62 from autophagic turnover. However, modified p62 remains autophagy-competent but is maintained as part of a broader TRIM21-protected p62 pool under IFN-stimulated conditions. Given that p62 ADPR is itself IFN-induced, preservation of this pool may allow p62-positive ADPR foci to contribute to degradation-independent functions.

### The PARP14-p62 axis regulates inflammatory gene expression in a context-dependent manner

The enrichment of innate immune regulators in the PARP14-dependent ADP-ribosylome and p62 interactome suggested that PARP14-dependent p62 foci may influence inflammatory signaling. We first tested this possibility in IFN-β-stimulated A549 cells by analyzing selected NF-κB, NRF2, and ISG target genes. p62 depletion or PARP14 inhibition did not substantially alter the expression of these genes after acute IFN-β stimulation (Fig. S4A and S4B). IFN-β stimulation alone may not fully capture the complexity of innate immune signaling. While IFN-β primarily activates the JAK-STAT pathway downstream of IFNAR, physiological inflammatory responses often involve simultaneous activation of multiple innate immune signaling axes, driven by IRF3/IRF7, NF-κB-, and IFN-dependent transcriptional programs^61–64^. Therefore, rather than testing multiple combinations of artificial stimuli in A549 cells, we sought to identify a cancer cell model with constitutive inflammatory signaling. Such basal inflammatory activity is frequently associated with a viral mimicry state, in which endogenous nucleic acids or retroelement-derived transcripts chronically activate innate immune pathways^65–67^. To identify suitable models, we used a previously established ISG score^16^ and calculated it for non-small cell lung cancer cell lines from the DepMap database (Fig. S4C). This analysis showed that A549 cells exhibit low basal inflammatory signaling, consistent with their low basal ADPR and their requirement for IFN stimulation. NCI-H1975 cells displayed an intermediate ISG score, which may explain why cysteine ADPR of p62 could already be detected in this cell line under unstimulated conditions. In contrast, NCI-H596 cells showed the highest ISG score among the analyzed models and were therefore selected for further experiments.

Consistent with the elevated basal ISG score, we detected low but detectable basal p62 ADPR in NCI-H596 cells upon endogenous p62 IP (Fig. 5A). We therefore investigated whether p62 and PARP14 contribute to the constitutive inflammatory gene expression observed in this model. To test this, we performed siRNA knockdown of p62 and PARP14 and also tested the contribution of PARP14 catalytic activity using the PARP14 inhibitor. Inflammatory gene expression was reduced across all conditions (Fig. 5B), suggesting that PARP14 supports basal inflammatory signaling in NCI-H596 cells. The similar effect observed after p62 depletion further indicates that this regulation may occur, at least in part, through a p62-dependent mechanism.

**Figure 5:**
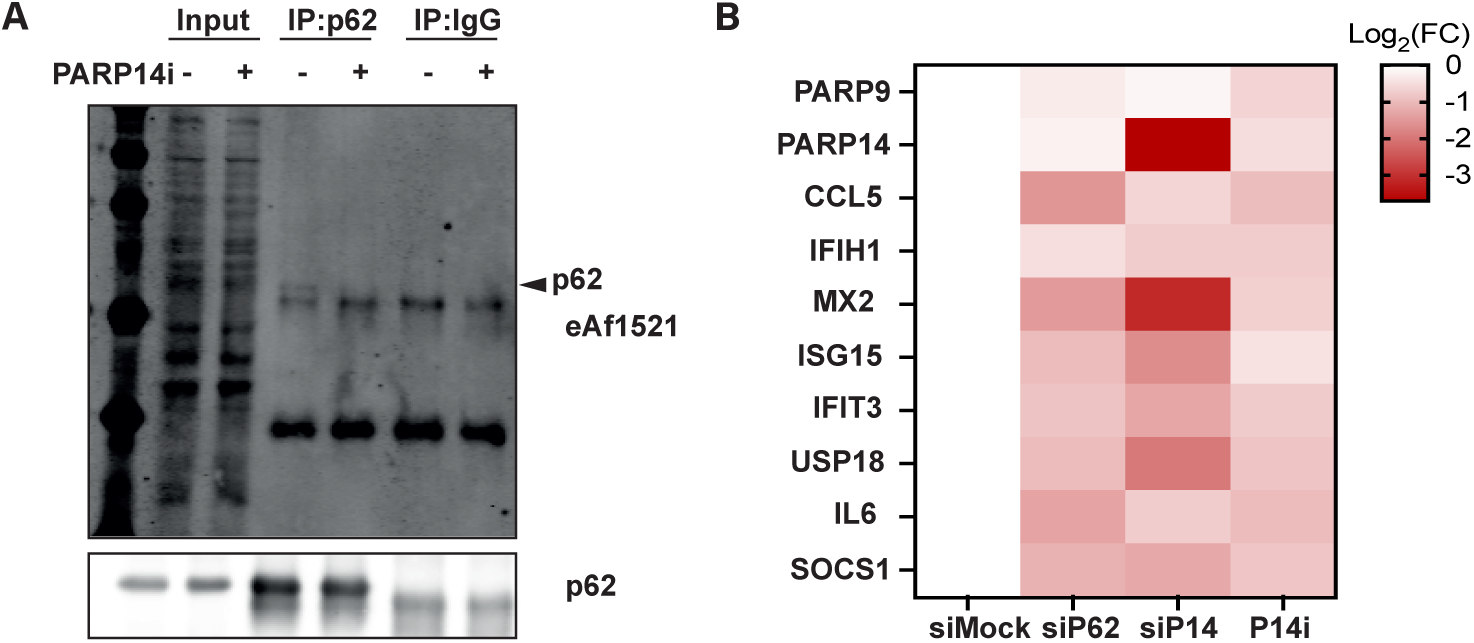
PARP14 and SQSTM1/p62 contribute to inflammatory signaling in a context- dependent manner. **(A)** Immunoblot analysis of endogenous p62 immunoprecipitation from NCI-H596 cells treated with PARP14i (P14i) (20 nM) for 24 h. **(B)** qPCR analysis of selected ISGs in NCI-H596 cells after treatment with 20 nM PARP14i (P14i) for 48 h or siRNA-mediated knockdown of p62 or PARP14 (siP14) for 48 h.

### SQSTM1/p62 interacts with PARP14-dependent ADP-ribosylation targets associated with the innate immune response

The observation that perturbation of either p62 or PARP14 reduced inflammatory gene expression in NCI-H596 cells suggested that both proteins may contribute to a shared inflammatory signaling network. However, the mechanistic link between p62 ADPR and this transcriptional phenotype remained unclear. We therefore asked whether p62 primarily acts as a PARP14 substrate or whether it also functions as a scaffold that organizes additional PARP14-dependent ADPR targets, thereby forming local ADPR hubs that could contribute to PARP14-dependent inflammatory signaling. To test this, we performed endogenous p62 immunoprecipitation followed by mass spectrometry (co-IP MS) in A549 cells treated with IFN-β alone or in combination with PARP14i. The overall identity of detected p62 interactors was largely preserved between conditions. However, the relative enrichment of individual interactors differed, indicating that PARP14 catalytic activity influences the composition or stability of specific p62-associated protein complexes (Fig. 4D). Among the proteins more strongly associated with p62 upon IFN treatment plus PARP14 inhibition were several IFN-induced ARTs as well as ubiquitin ligases, including RNF114 and RNF166.

Consistent with the idea that p62 foci represent ADPR hubs, 30% of proteins detected in the PARP14-dependent ADP-ribosylome were also present in the IFN-β-dependent p62 interactome (Fig. S3C). In addition, immunoblot quantification showed that p62 knockdown reduced the overall IFN-β- induced ADPR signal by approximately 25% (Fig. 4E). This suggests that p62 contributes to efficient IFN-induced ADPR of a subset of ADPR targets, consistent with a scaffolding function of p62 foci. Functional enrichment of the p62 interactome did not reveal a dominant enrichment of canonical autophagy pathways. Instead, enriched terms were associated with mitochondrial organization and innate immune regulation (Fig. S3D). Comparison of the IFN-β and IFN-β+PARP14i conditions further suggested that a subset of p62-associated proteins may depend on PARP14 activity for efficient recruitment. Proteins more strongly enriched under IFN-β treatment included components of the TRiC/CCT chaperonin complex and several proteasomal subunits (Fig. S3E and S3F). Together, these data support a model in which p62 foci function as scaffolds for PARP14-dependent ADPR of multiple immune- and proteostasis-associated proteins.

## DISCUSSION

In this study, we identify the selective autophagy receptor SQSTM1/p62 as a major ADPR target in lung epithelial cancer cell lines. In A549 cells, p62 ADPR was low under basal conditions but was induced by multiple inflammatory stimuli and dependent on the catalytic activity of PARP14. Site-resolved analysis showed that p62 is modified on several acceptor residues, with cysteine ADPR representing the most prominent and reproducible site. Consistent with this, the broader PARP14- dependent ADPR landscape was enriched for cysteine acceptor sites and included proteins linked to innate immune signaling, ubiquitin regulation, and RNA restriction. We further show that PARP14- dependent ADPR is spatially organized within cytoplasmic p62 foci that are distinct from classical autophagosomes. Proteomic analysis revealed that several PARP14-dependent ADPR targets associate with p62, supporting a model in which p62-positive foci act as scaffolds for a subset of IFN-induced ADPR events. Finally, although ADP-ribosylated p62 remains competent for autophagic turnover, TRIM21 limits degradation of this modified p62 pool, suggesting that p62-associated ADPR foci may be preserved for degradation-independent functions with potential relevance to inflammatory signaling.

We identified cysteine ADPR as a prominent PARP14-dependent modification site, based on our PARP14-dependent ADP-ribosylome analysis and mutagenesis of p62 acceptor sites, which resulted in the loss of modification without affecting the physiological behavior of p62 (localization and ability to form foci) (Fig. S2F). In contrast to our results, the current literature suggests that PARP14 typically modifies substrates at acidic amino acid residues, such as glutamic and aspartic acid^8,68^. However, these previous studies were either based on *in vitro* modification of recombinant proteins or have been questioned due to potential confounding factors, such as nearby modifications by different posttranslational modifications^69^. Despite the strong evidence for PARP14-dependent modification of cysteine residues in this study, we cannot entirely rule out an indirect mechanism. PARP14 may act upstream of other ARTs, regulating their activity, localization, or substrate accessibility. This could enable an intermediate ART to modify target proteins on cysteine residues. This possibility is supported by our p62 interactome analysis, in which several ARTs were in complex with PARP14 (Fig. 4D). Therefore, while our data establish PARP14 as a major upstream regulator of cysteine ADPR under IFNβ-stimulated conditions, we cannot rule out PARP14-dependent regulation of downstream ARTs.

We further show that PARP14-dependent ADPR is spatially concentrated within p62 foci. The formation of p62 foci is well documented in the literature and is typically associated with autophagic turnover, as p62 aggregation is a prerequisite for this process^31,32^. However, these foci stained negative for the autophagosome marker LC3, arguing against their identity as canonical LC3-positive autophagosomes. This is consistent with recent reports describing similar structures as interferon-induced cytoplasmic ADP-ribose bodies (ICABs), highlighting that these IFN-induced ADPR foci represent a distinct biological entity^70^. PARP14 inhibition reduced the ADPR signal and PARP14 enrichment at these foci but did not abolish p62 foci or ubiquitin accumulation. Thus, PARP14 does not appear to initiate p62 body formation but uses it as a pre-existing platform to promote ADPR. This hypothesis is supported by both the overlap between the PARP14-dependent ADP-ribosylome and the p62 interactome and the reduction in overall ADPR following p62 depletion. Together, these findings support a model in which p62 acts not only as a substrate but also as a scaffold that concentrates PARP14-dependent ADPR within ubiquitin-rich cytoplasmic foci.

The link between PARP14-dependent ADPR and ubiquitin signaling is further supported by the identification of RNF31/HOIP, SPATA2, RNF114, and RNF166 as PARP14-dependent targets. These factors connect the dataset to LUBAC-dependent NF-κB signaling, ubiquitin-ADP-ribose hybrid mark biology and proteasome-dependent degradation. However, ubiquitin and M1-linked ubiquitin accumulation within p62 foci were not visibly reduced by PARP14 inhibition, suggesting that ubiquitin assembly occurs independently of PARP14 activity. PARP14 may therefore modify proteins within ubiquitin-rich p62 foci rather than controlling formation of the ubiquitin assemblies themselves. The presence of RNF114 and RNF166 in this network is particularly interesting, as it suggests that p62- positive ADPR foci may also serve as sites for generating ubiquitin-ADP-ribose hybrid marks, a process previously linked to proteasomal degradation^71^.

Given the established role of p62 in selective autophagy^25,30,72^, we initially considered that p62 ADPR near functionally important cysteine residues could regulate p62 autophagic activity. However, PARP14-dependent p62 ADPR did not detectably alter p62 foci formation, nucleocytoplasmic shuttling, or LC3 binding. ADP-ribosylated p62 accumulated after chloroquine treatment and was recovered in LC3-EGFP pulldowns, indicating that the modified pool remains competent for autophagic turnover. Moreover, PARP14 knockdown did not phenocopy p62 depletion in the degradation of Hepatitis C core protein, suggesting that this specific p62-dependent cargo turnover does not require PARP14-dependent p62 ADPR.

Instead, our data suggest that under IFN-induced conditions, p62 is protected from autophagic degradation. TRIM21 depletion reduced both total and ADP-ribosylated p62, and this effect was rescued by chloroquine. Thus, TRIM21 does not appear to selectively protect only the ADP-ribosylated form of p62, but rather limits autophagic turnover of p62 more broadly under IFN-stimulated conditions. Importantly, the ADP-ribosylated fraction remains sensitive to this regulation, indicating that modified p62 is autophagy-competent but maintained as part of a TRIM21-protected p62 pool. This supports a model in which p62-associated ADPR foci are maintained for degradation-independent functions, potentially linked to innate immune regulation and protein quality control.

p62 ADPR was sensitive to SARS-CoV-2 nsP3 Mac1 activity both in cells and in the reconstituted *in vitro* assay (Fig. S1H-J). However, these data do not establish p62 as a direct viral macrodomain substrate. *In vitro*, Mac1 also strongly reduced PARP14 auto-ADPR, suggesting that the loss of p62 trans-modification may be a consequence of ADPR hydrolysis of PARP14. This interpretation is supported by our site-resolved data showing that PARP14 itself is extensively ADP-ribosylated on acidic residues, which are established substrates for several macrodomains^73^. Nevertheless, these findings indicate that PARP14-dependent ADPR is sensitive to viral macrodomain activity and provides a potential link between PARP14-dependent ADPR and host-virus conflict.

The functional consequences of the PARP14-p62 axis appear to be context-dependent. In A549 cells, p62 depletion or PARP14 inhibition did not substantially alter selected NF-κB, NRF2, or ISG target genes after acute IFN-β stimulation. In contrast, depletion of p62 and PARP14, and PARP14 inhibition, reduced inflammatory gene expression in the constitutively inflammatory NCI-H596 model. These findings suggest that the PARP14-p62 axis may become functionally relevant in cellular states with chronic or multilayered innate immune activation. However, additional experiments will be required to determine whether this phenotype depends specifically on p62 ADPR or on independent functions of PARP14 and p62 within the same inflammatory signaling network.

In summary, our findings identify p62 as a major IFN-induced, PARP14-dependent ADPR target and reveal that PARP14-dependent ADPR is concentrated within p62 foci. Rather than acting solely as an autophagy receptor, p62 appears to function as a scaffold for ADP-ribosylated immune- and proteostasis-associated protein assemblies. Together, these data support a model in which PARP14- dependent cysteine-enriched ADPR marks p62 foci and connects interferon signaling to spatially organized innate immune regulation.

## DATA AVAILIBILITY

Original data is available upon request. The mass spectrometry proteomics data have been deposited to the MassIVE repository with the dataset identifier PXD082766.

## ACKNOWLEDGMENTS

We thank Sofia Andretta and Lukas Muskalla (University of Zurich) for providing editorial assistance. We thank the Center for Microscopy and Image Analysis (ZMB) and the Functional Genomics Center Zurich (FGCZ) for their services and assistance. We further thank Jonas Grossmann (FGCZ) for methodological recommendations and bioinformatics support. This work was supported by the Swiss National Science Foundation [31003A_176177 and 310030_205202 to M.O.H.].

## AUTHOR CONTRIBUTIONS

**Project conceptualization and administration:** D.K. and M.O.H. (lead)

**Data curation and Formal analysis:** D.K. (lead), D.M.L.P., and S.S. (lead mass spectrometry)

**Investigation (specific experiments)**: D.K. (lead), S.S., and D.N. (supporting)

**Visualization and validation**: D.K. (lead)

**Methodology:** D.K. (lead), D.M.L.P., and S.S. (lead mass spectrometry)

**Writing, review & editing of MS**: D.K. and M.O.H. (lead), D.M.L.P. (supporting)

**Declaration of interests:** The authors declare no conflict of interest.

## MATERIALS and METHODS

### Cell Culture

Wildtype A549 cells were grown in high-glucose Dulbecco’s modified Eagle’s medium (DMEM) supplemented with 10% (v/v) fetal calf serum (FCS) and 5% (v/v) penicillin/streptomycin (P/S). Genetically engineered cell lines (A549 p62-HA, A549 p62-HA C113A, C289A, C290A, C331A mutant, A549 LC3-EGFP, and A549 nsP3 Mac1) were grown in high-glucose DMEM medium supplemented with 10% FCS, 5% P/S, and 30 μg/mL Blasticidin (Invivogen). Transgene expression of genetically engineered cell lines was induced using doxycycline. Treatment time and concentration are indicated in the figure legends. All cell lines were cultured at 37°C in a humidified atmosphere containing 5% CO_2_.

### Generation of genetically engineered cell lines

Inducible overexpression cell lines were generated by transposase-mediated integration. Therefore, the coding sequences of p62-HA (Addgene #28027), LC3-EGFP (Addgene #11546), and SARS-CoV-2 nsP3 Mac1 (Kind gift by Sarah Knapp, RWTH Aachen) were subcloned into a Sleeping Beauty vector (Addgene #60506). The Sleeping Beauty vector was co-transfected with Sleeping Beauty Transposase (Addgene #34879) using Lipofectamine 3000 (Thermo Fisher). 72 h post-transfection, cells were subjected to Blasticidin selection. ADPR-deficient p62-HA mutant (C113A, C289A, C290A, C331A) was generated by standard site-directed mutagenesis PCR.

### Drug treatment

PARP14 was inhibited using 20 nM of RBN012759 (PARP14i) (MedChemExpress LLC, Monmouth Junction, NJ). If not stated otherwise, RBN012759 was co-treated with the following p62-ADPr inducing agents at the following concentrations by direct addition to the culture medium IFN-β (1000U/mL, Peprotech), IFNγ (100U/mL, Peprotech). The p62-ADPr inducing agent Poly(I:C) LMW (Invivogen) was transfected using Lipofectamine3000 at a concentration of 10 ng/mL. Treatment durations are indicated in the figure legends.

### siRNA transfection

siRNA-mediated knockdowns were performed via reverse transfection using Lipofectamine RNAiMAX (Thermo Fisher). 10 nM of siRNA were mixed with 3 μL lipofectamine in 500 μL serum-free OptiMEM and incubated for 20 min at room temperature (RT). The transfection mix was added dropwise to the cells, which were then incubated for 48 h. A non-targeting siRNA (siMOCK) was used as a control for each experiment. siRNA sequences are listed in Table 2.

### Immunoblotting

Cells were lysed with LDS lysis buffer (60 mM Tris-HCl pH 6.5, 2% LDS, and 10% Glycerol) and denatured at 70°C for 3 min. The samples were sonicated, and proteins were separated on 10% Bis-Tris gels at 100V. A wet transfer was performed onto PVDF membranes at 100 V for 1.5 h at room temperature (RT), and the membranes were blocked with 5% non-fat milk in TBS-T for 1h at RT. Primary antibodies were diluted in 5% non-fat milk in TBS-T and incubated with the membrane at 4°C overnight. Membranes were then washed three times in TBS-T before being incubated with a secondary antibody diluted in TBS-T for 1 h at RT. After three washes in TBS-T, specific proteins/bands were detected with the Odyssey infrared imaging system (LI-COR). The following primary and secondary antibodies were used at the indicated dilutions: rabbit anti-SQSTM1/p62 (ab155686, Abcam, 1:1000), mouse anti-SQSTM1/p62 (ab56416, Abcam, 1:1000), mouse anti-Tubulin (T6199, Sigma, 1:5000), rabbit anti-PARP9 (ab53796, Abcam, 1:250), rabbit anti-PARP14 (ab224352, Abcam, 1:500), rabbit anti-TRIM21 (12108-1-AP, Proteintech), rabbit anti-HA (ab9110, Abcam, 1:5000), mouse anti-FLAG (F3165, Sigma, 1:500), mouse anti-GFP (66002-1-Ig, Proteintech, 1:20000), mouse eAf1521 macrodomain (Hottiger laboratory, final concentration 0.4 ng/µL), IRDye 800 CW goat anti-rabbit IgG (LI-COR, 1:10000) and IRDye 680RD goat anti-mouse IgG (LI-COR, 1:10000).

### Immunofluorescence (IF)

Cells were grown on glass coverslips before the treatments. Following the treatments, cells were fixed with 4% formaldehyde in PBS for 20min at RT and permeabilized with 0.2% Triton-X-100 in PBS for 10min. DMEM supplemented with FCS and 0.1% Triton-X-100 was used as a blocking solution to block the cells for 1 h at RT. Next, coverslips were incubated in the primary antibody diluted in blocking buffer at 4°C overnight. Coverslips were washed three times with PBS and incubated with a secondary antibody diluted in a blocking buffer for 1h at RT. After three washes, coverslips were stained with a 0.1µg/mL DAPI solution in PBS for 20min at RT. Coverslips were again washed three times with PBS and mounted on microscope slides using Mowiol® 4-88. Images were acquired on an Olympus ScanR HCS and a Leica Thunder microscope. Contrast and brightness adjustments were performed in Fiji (ImageJ)^75^, and the same settings were applied to the entire image set within a single experiment. Quantifications of immunofluorescence colocalization and mean fluorescence intensity were performed with a custom CellProfiler^76^ image analysis pipeline. The following primary and secondary antibodies were used for IF: rabbit anti-SQSTM1/p62 (ab155686, Abcam, 1:250), mouse anti-SQSTM1/p62 (ab56416, Abcam, 1:250), mouse anti-HA (901502, Biolegend, 1:500), rabbit anti-HA (ab9110, Abcam, 1:250), rabbit anti-PARP14 (ab 229756, Abcam 1:250), Mono- and polyubiquitinylated conjugates FK2 (BML-PW8810-0100, Enzo Life Sciences, 1:200), rabbit anti-Calnexin (2679, CellSignaling, 1:100), rabbit anti-M1 Ubiquitin (ZRB2114, Sigma, 1:1000), goat anti-rabbit IgG(H+L) Alexa Fluor 488 (A11008, Invitrogen, 1:250), goat anti-mouse IgG (H+L) Cy3 (115-165-003, Jackson Immunoresearch, 1:250).

### Quantitative real-time PCR (qPCR)

Isolation of RNA was performed with the NucleoSpin RNA II kit (Macherey-Nagel). RNA concentrations were determined by Nanodrop, and RNA was reverse transcribed using the High-Capacity cDNA Reverse Transcription Kit (Thermo Fisher/Applied Biosystems) according to the manufacturer’s instructions. cDNA was used for qPCR using the KAPA SYBR® FAST (Sigma Aldrich) master mix on a Rotor-Gene Q 2plex HRM system (Qiagen) or QuantStudio5 System (Applied Biosystems). qPCR data were normalized to RPS12, and ΔΔCt analysis was performed relative to the control sample. Primers are listed in Table 1.

**Table 1:**
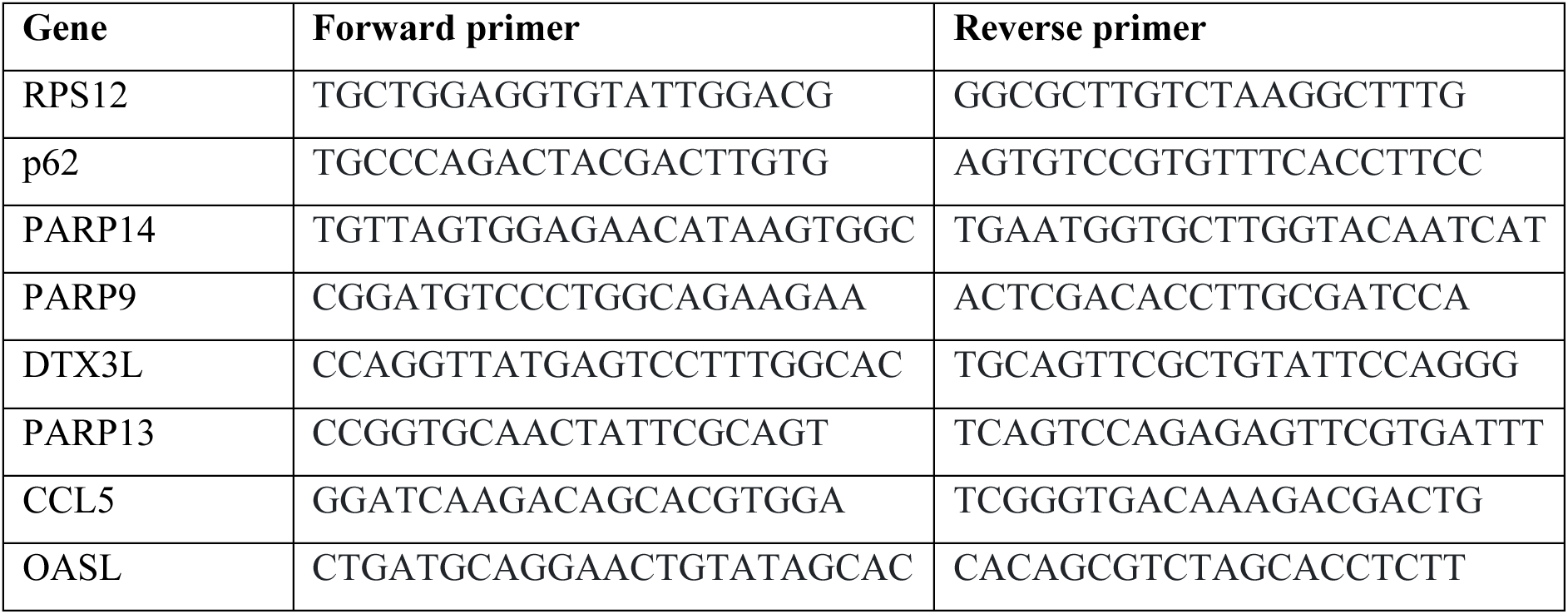
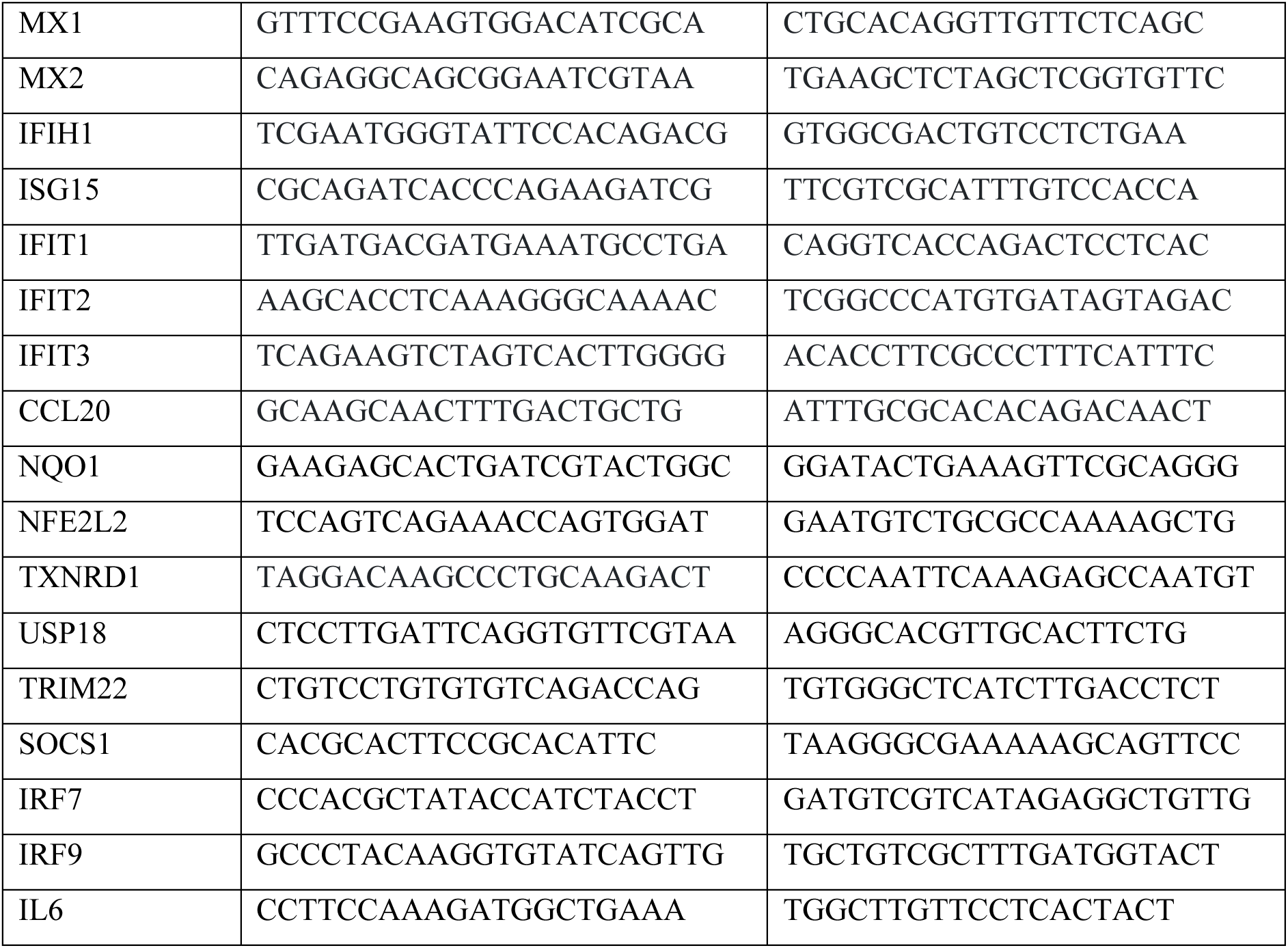
List of primers.

**Table 2:**
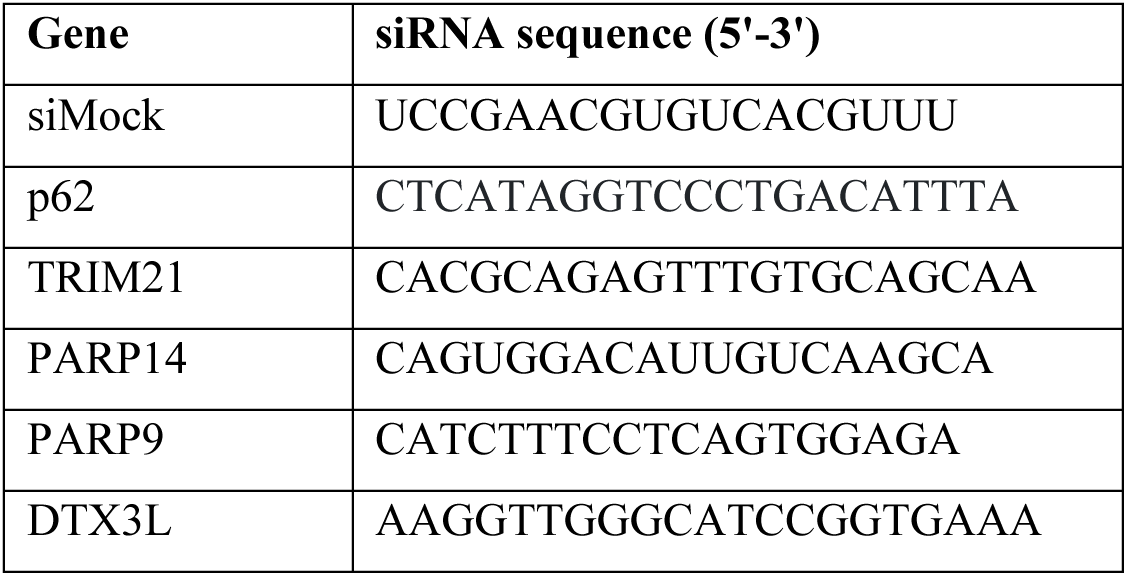
List of siRNA.

### Immunoprecipitation (IP)

Cells were lysed with non-denaturing lysis buffer to maintain protein-protein interactions (10 mM Tris-HCL pH 7.5, 100mM NaCl, 0.5 mM EDTA, 0.5% NP-40, 40 μM PJ-34, 1 μg Leupeptin, 1 μg Pepstatin, 1 μg Bestatin, 100 μM PMSF) at 4°C for 30 min on a head-over-tail rotor. Samples were subsequently centrifuged at 14000g for 15min at 4°C, and the supernatant was incubated with either 1μg of rabbit anti-SQSTM1/p62 antibody (ab155686, Abcam) or rabbit IgG antibody (PP64B, Millipore) as a control for 2 h at 4°C. Protein A Dynabeads (Invitrogen) were equilibrated with wash buffer (NP-40-free lysis buffer), and 30 μL of beads were subsequently added to the lysates and incubated for 1h at 4°C. Beads were then washed three times with the wash buffer. 30 μL of 2x LDS lysis buffer was added to the samples, which were then boiled for 3min at 70°C and analyzed by immunoblotting.

### GFP-Trap Immunoprecipitation

Cells were harvested and processed as described for our standard immunoprecipitation (IP) protocol. For immunoprecipitation of GFP-tagged proteins, 25 μL of GFP-Trap® Magnetic Particles M-270 (gtd, ChromoTek) were incubated with the lysate for 1 h at 4°C. Samples were then again processed as previously described.

### ADPr-peptide enrichment

For ADP-ribosylome analysis, A549 cells were treated with IFN-β (100 U/mL) and 20nM RBN012759 (PARP14i) for 24 h. Cells were washed twice with PBS, trypsinized for 5min at 37°C, and collected by centrifugation at 300g for 5min. For each condition (n=4/condition), two cell pellets were processed for protease digestion. One cell pellet was lysed in GndHCl lysis buffer (6 M GndHCl, 50 mM Tris pH 8.0), sonicated, and stored at −80°C until LC MS-MS analysis. The other cell pellet was lysed in urea lysis buffer (8 M urea, 20 mM HEPES pH 7.0, 5 mM DTT, 5 mM MgCl_2_ with 10 µM olaparib, 1 µM ADP-HPD), 750U benzonase was added to each sample and incubated at room temperature for 10min with rotation and for an additional 5 min at 37°C with agitation (220 RPM). Stored at −80°C until LC-MS/MS analysis. An aliquot corresponding to 10 mg of protein was taken for both the GndHCl and Urea lysates. Protein disulfide bridges were reduced with 5mM Tris(2-carboxyethyl) phosphine (TCEP) and alkylated with 10 mM 2-Chloroacetamide (CAA) in the dark at 30°C for 30 min. The GndHCl lysates were diluted to a final concentration of 0.5M GndHCl in PARG buffer (50 mM Tris-HCl pH 8.0, 50 mM NaCl, 10 mM MgCl_2_, 250 µM DTT) and digested with modified Porcine Trypsin (1:25; Sigma) for 3 hours at 25°C. At the same time, the Urea lysates were diluted to a final concentration of 0.5M Urea in digestion buffer (20mM sodium acetate, pH 5.0, 5 mM DTT), Arg-C Ultra was added at a 1:2000 ratio (protease:protein w/w) and incubated for 3 h at 37°C. The digests were then pooled and acidified to pH 4.0. ADPr-peptide enrichments were carried out as previously described^33,77^ with the following protocol modifications. PARG-mediated PAR-to-MAR peptide ADPr-modification reduction was not carried out, and the peptides were enriched using Af1521-WT (0.5 mL beads/20 mg lysate) and eAF1521 (1.0 mL beads/20 mg lysate) macrodomain affinity enrichment for 2 h at 4°C. The enriched samples were then prepared for MS analysis as described previously^33^.

### Liquid chromatography and mass spectrometry analysis

Identification of ADP-ribosylated peptides from untreated and treated cells was performed on an Orbitrap Lumos mass spectrometer (Thermo Fisher Scientific) coupled to an ACQUITY M class UPLC liquid chromatograph (Waters). We applied an ADPr product-dependent analysis called HCD-PP- EThcD^78^. Solvent compositions in channels A and B were 0.1% formic acid in water and 0.1% formic acid in acetonitrile, respectively. Peptides were loaded onto a nanoEase M/Z Symmetry (Waters) trap column, 18 μm × 20 mm, packed with C18 material, 5 μm, 100 Å, and separated on an analytical nanoEase M/Z HSS T3 Column (Waters, 75μm × 200mm) packed with reverse-phase C18 material (Waters, 1.8μm, 100 Å). Peptides were eluted over 110min at a 300 nl/min flow rate. A linear elution gradient protocol was used from 3 to 25% B for 95 min, followed by 35% B for 5 min, and a wash step at 95% B for 5 min, respectively. Full-scan MS spectra (350−2000 m/z) were acquired at a resolution of 120,000, with an AGC target set at 4e5 and a maximum injection time of 50 ms. High-energy HCD MS/MS spectra (collision energy at 38%) were acquired at a resolution of 30,000, with the AGC target set at 5e4 and a maximum injection time of 60 ms. Two or more observed ADPr fragment peaks (136.0623, 250.0940, 348.07091, and 428.0372) in the high-energy data dependent HCD scan triggered additional high-quality HCD and EThcD MS/MS scans (resolution of 120,000, AGC target set at 5e5, injection time of 240ms, collision energy for HCD scan at 35%).

### ADP-ribosylome Data Analysis

For LFQ based comparisons between conditions, raw MS data were searched against the reviewed human UniProt proteome including reversed decoy sequences using MSFragger within FragPipe (v22.0). Searches were performed in DDA mode with a precursor mass tolerance of ±20 ppm with mass calibration and parameter optimization enabled and a fragment ion tolerance of 10ppm, matching b, y, c, and z product ions. Peptides were generated using a combined tryptic/Arg-C digestion strategy (cleavage C-terminal to K/R and to R, respectively, with proline exclusion for Arg-C), requiring full enzymatic specificity for both enzymes and allowing up to 2 missed cleavages per enzyme, with peptide lengths restricted to 7-50 residues (500-5000 Da).

To detect ADPR, searches were performed in FragPipe’s labile search mode. A 0 Da mass offset for unmodified window and a +541.0611 Da offset corresponding to ADPr was applied to peptides containing candidate acceptor residues (Cys, Asp, Glu, His, Lys, Arg, Ser, Thr, Tyr). Because cysteine was also subject to fixed carbamidomethylation (+57.02146 Da), a second, cysteine-specific offset of +484.03964 Da (541.0611 - 57.02146) was included to capture ADP-ribosylated cysteine residues that did not additionally carry the alkylation adduct. Spectra were required to contain diagnostic ADPr fragment ions (m/z 136.06232, 250.09401, 348.07036, 428.03669, and 584.09018) at a relative intensity exceeding 0.03 of the base peak, and delta-mass localization was enabled to assign modification sites within matched peptides. Carbamidomethylation of cysteine (+57.02146 Da) was set as a fixed modification; oxidation of methionine (+15.9949 Da), N-terminal protein acetylation (+42.0106 Da), and ADPR (up to 2 per peptide, restricted to the residues above) were included as variable modifications, with a maximum of 3 total variable modifications allowed per peptide. Site-level intensity data were analyzed using the prolfqua R package from the FGCZ^79^.

For determination of ADPr site localization, MS and MS/MS spectra were converted to Mascot generic format (MGF) using Proteome Discoverer, v2.1 (Thermo Fisher Scientific, Bremen, Germany). Separate MGF files were created from the raw file for each type of fragmentation (HCD and EThcD) using a dedicated rule in the converter control^80^. Mascot searches were carried out as previously described^81^. A peptide tolerance of 8 ppm and MS/MS tolerance of 0.03 Da were used. Enzyme specificity was set to trypsin, allowing up to 4 missed cleavages. The MGFs were searched against the target-decoy UniProtKB human database (taxonomy 9606, canonical sequences and reviewed entries only, downloaded on 2019/07/06). N-terminal protein acetylation was set as a variable modification. C, D, E, H, K, R, S, T and Y residues were set as variable ADPr acceptor sites with a mass shift of 541.0611 Da. Carbamidomethylation was set as a fixed modification on C and oxidation as a variable modification on M. The neutral losses from the ADPr 249.0862 Da, 347.0631 Da, and 583.0829 Da were scored in HCD fragment ion spectra. For HCD and EThcD fragment ion spectra, the marker ions at *m/z* 428.0372, 348.0709, 250.0940, 136.0623 were ignored for scoring. Peptides were considered as correctly identified when a Mascot score >15 and an expectation value <0.05 were obtained. For the ADPR site analyses, peptides identified with a localization score >95%, were used.

### Co-immunoprecipitation and proteome trypsin digestion

Following SQSTM1/p62 immunoprecipitation using the above protocol, the proteins were subjected to trypsin digestion using an on-bead digestion protocol. Briefly, the beads were washed twice with digestion buffer (50 mM AmBic/2 mM CaCl_2_, pH 8.2) and then resuspended in digestion buffer containing Sequencing Grade Trypsin (10 ng/µL; Promega). Beads were incubated with agitation overnight at 37°C. Peptides were collected, and the beads were washed once with the digestion buffer. Samples were pooled and then dried in a SpeedVac. They were then dissolved in 3% ACN with 0.1% formic acid and stored at −20°C until MS analysis.

The 6 M GndHCl protein lysates from the ADP-ribosylome analyses were used for whole proteome LFQ analyses. Protein disulfide bridges were reduced with 5 mM Tris(2-carboxyethyl)phosphine (TCEP) and alkylated with 10 mM 2-Chloroacetamide (CAA) in the dark at 95°C for 10 min. 40 µg of proteins were prepared for digestion using the filter-aided sample preparation (FASP) methodology^82^ and digested with Sequencing Grade Trypsin (1:20; Promega) overnight at 37°C. The samples were then acidified with TFA and salts removed using C18 spin columns (Thermo Scientific). The peptides were eluted with 40 µl of 60% ACN, 0.1% TFA, dried to completion, and then re-dissolved in 3% ACN, 0.1% formic acid to a final peptide concentration of 0.5 µg/µl.

### Liquid Chromatography and Mass Spectrometry Analysis

LC-MS/MS analyses were performed on an Orbitrap Exploris mass spectrometer (Thermo Fisher Scientific), coupled to ACQUITY UPLC liquid chromatographs (Waters). Peptides were loaded on a commercial MZ Symmetry C18 Trap Column (100Å, 5 µm, 180 µm x 20 mm, Waters) followed by nanoEase MZ C18 HSS T3 Column (100Å, 1.8 µm, 75 µm x 250 mm, Waters). Peptides were eluted over 110 min at a flow rate of 300 nL/min. An elution gradient protocol from 2% to 25% B, followed by two steps at 35% B for 5 min and at 95% B for 5 min, respectively, was used. The mass spectrometer was operated in data-dependent mode (DDA), acquiring a full-scan MS spectra (300−1’800 m/z) at a resolution of 120’000 at 200 m/z after accumulation to a target value of 500’000. Data-dependent MS/MS were recorded in the linear ion trap using quadrupole isolation with a window of 0.8 Da and HCD fragmentation with 30% fragmentation energy. The ion trap was operated in rapid scan mode with a target value of 10’000 and a maximum injection time of 50 ms. Only precursors with intensities above 5’000 were selected for MS/MS, and the maximum cycle time was set to 3 s. Charge state screening was enabled. Singly, unassigned, and charge states higher than seven were rejected. Precursor masses previously selected for MS/MS measurement were excluded from further selection for 20 s, and the exclusion window was set at 10 ppm. The samples were acquired using internal lock-mass calibration at m/z 371.1012 and 445.1200.

### Protein identification and label-free protein quantification

The acquired raw MS data were processed using MaxQuant (version 1.6.2.3), followed by protein identification using the integrated Andromeda search engine^83^. Spectra were searched against the Swiss- Prot human reference proteome (taxonomy 9606, version from 2019-07-09), concatenated with its reversed-decoy fasta database and common protein contaminants. Methionine oxidation and N-terminal protein acetylation were set as variables. Enzyme specificity was set to trypsin/P, allowing a minimum peptide length of 7 amino acids and a maximum of two missed cleavages. MaxQuant Orbitrap default search settings were used. The maximum false discovery rate (FDR) was set to 0.01 for peptides and 0.05 for proteins. Label-free quantification was enabled, and a 2 min window for matching between runs was applied. In the MaxQuant experimental design template, each file is kept separate in the experimental design to obtain individual quantitative values. Protein fold changes were computed based on Intensity values reported in the proteinGroups.txt file. A set of functions implemented in the R package SRMService ^84^was used to filter for proteins with 2 or more peptides, allow up to 4 missing values, normalize the data using a modified robust z-score transformation, and compute p-values using a t-test with pooled variance.

### *In vitro* ADP-ribosylation assay

*In vitro* assays were performed in a 50 μl reaction volume using 0.5μg of recombinant full-length PARP14-His (Kind gift by Karla Feijs-Žaja, RWTH, Aachen) and recombinant full-length p62-GST (abcam, ab132366). Recombinant SARS-CoV-2 Mac-His was a kind gift from Aikaterini Tsika (University of Patras). Reactions were performed in reaction buffer (20 mM HEPES, pH 7.0, 150 mM NaCl, 1mM DTT, 1mM MgCl_2_) with the addition of 40 μM NAD^+^ and incubated for 1h at 30°C on a shaker at 300 rpm. Reactions were stopped by adding 1x LDS loading buffer.

### Statistical Analysis

Fluorescence intensity measurements were normalized, and the means of three independent experiments were compared. Pearson correlation was again calculated from the means of three independent experiments for statistical analysis. For all experiments with two groups, p values were determined by Student’s t-test with *, p<0.05; **, p <0.005; ***, p < 0.0005, **** p < 0.0001. For experiments with three or more groups, p-values were determined using one-way ANOVA with multiple comparisons. The exact statistical test is detailed in the figure legends.

---

**Figure S1:**
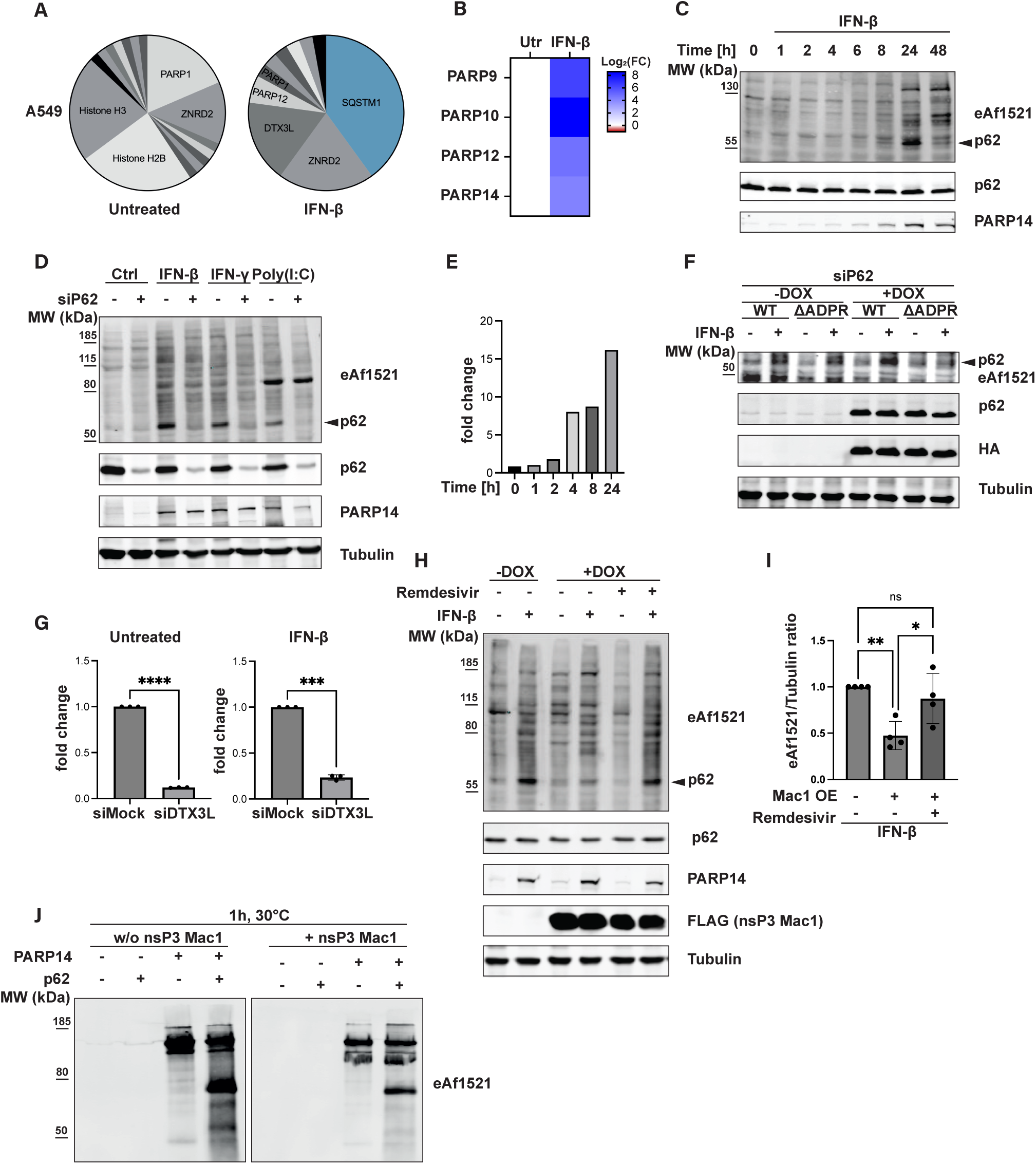
SQSTM1/p62 is ADP-ribosylated at cysteine residues in a PARP14 dependent manner. **(A)** Pie chart showing unique ADP-ribosylated proteins identified by LC-MS/MS in A549 cells treated for 24h with IFN-β (1000 U/mL), in comparison to untreated control cells. **(B)** Heatmap showing qPCR analysis in A549 cells treated with IFN-β (1000 U/mL) for 24 h. **(C)** Immunoblot analysis of A549 cells treated with IFN-β (1000U/mL) for up to 48 h. Cells were harvested at the indicated time points post IFN-β treatment. **(D)** Immunoblot of A549 cells after p62 knockdown for 48h. For the last 24h of the experiment, cells were either treated with IFN-β (1000 U/mL) or IFNγ (100 U/mL). Poly(I:C) (10 ng/mL) was transfected in the last 8 h of the experiment. **(E)** qPCR analysis of *PARP14* A549 cells treated with IFN-β (1000U/mL) for up to 24 h. Cells were harvested and analyzed at the indicated time points. **(F)** Immunoblot of stable A549 p62-HA Wildtype (WT) and A549 p62-HA ADPr-deficient mutant (ΔADPR) cell lines after knockdown of endogenous p62 for 48 h. The ΔADPr mutant contains the following mutations: C113A, C289A, C290A, and C331A. Cells were treated with IFN-β (1000U/mL) for the last 24h. Expression of the respective p62-HA construct was induced with 1μg/mL doxycycline for the last 6h of the experiment. **(G)** Knockdown of DTX3L, shown in Fig. 1E, was confirmed by qPCR analysis. P values were determined by a paired, two-tailed Student’s t-test. **(H)** Immunoblots of A549 cells with stable overexpression of a doxycycline-inducible FLAG-tagged SARS-CoV-2 nsP3 macrodomain 1 construct for 24 h using 500 ng/mL of doxycycline. Cells were co- treated with IFN-β (1000 U/mL) and remdesivir (20 μM), a prodrug whose active metabolite has been reported to inhibit Mac1 activity^74^. **(I)** Quantification of the western blot shown in (H). **(J)** *In vitro* assay with recombinant full-length PARP14 and full-length p62 in the presence and absence of recombinant SARS-CoV-2 Mac1 (nsP3 Mac1). The reaction was performed for 1 h at 30°C using 40 μM NAD^+^.

**Figure S2:**
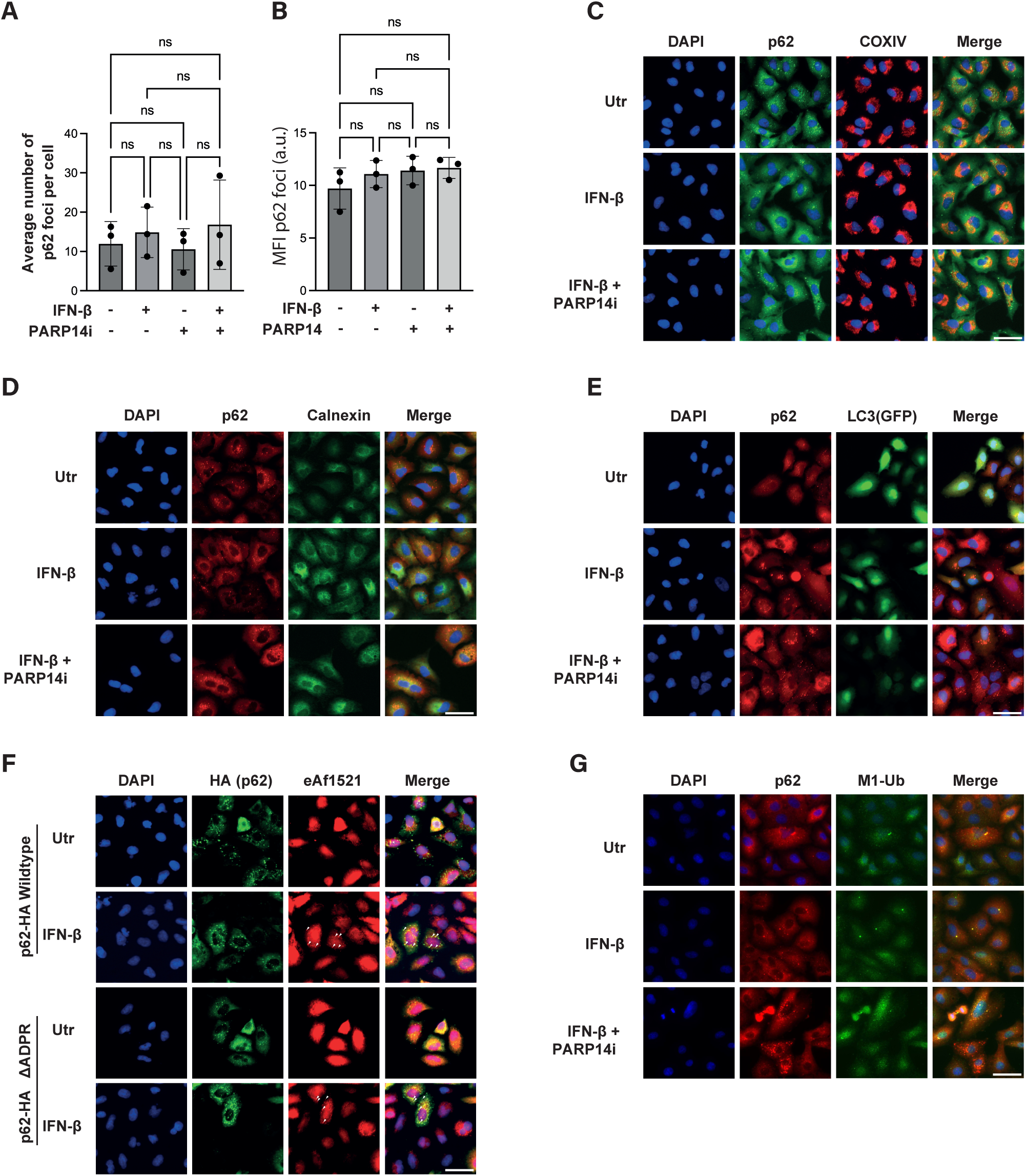
SQSTM1/p62 forms cytoplasmic foci which colocalize with ADP-ribosylation, PARP14 and Ubiquitin. **(A/B)** Quantification of p62 foci number **(A)** and mean fluorescence intensity (MFI) of p62 foci **(B)** from the data sets shown in Fig. 2A. **(C/D)** Immunofluorescence images of A549 cells treated with IFN- β (1000 U/mL) and PARP14i (20 nM) for 24 h. **(E)** A549 LC3-EGFP cells were treated with 500 ng/mL doxycycline for 24h to induce LC3-EGFP expression. Cells were additionally treated with IFN-β (1000 U/mL) and PARP14i (20 nM) for 24 h. **(F)** Stable A549 p62-HA Wildtype (WT) and A549 p62-HA ADPr-deficient mutant (ΔADPR) cell lines after knockdown of endogenous p62 for 48 h. The ΔADPr mutant contains the following mutations: C113A, C289A, C290A and C331A. Cells were treated with IFN-β (1000 U/mL) for the last 24h. Expression of the respective p62-HA construct was induced with 1 μg/mL doxycycline for the last 4h of the experiment. **(G)** Immunofluorescence images of A549 cells treated with IFN-β (1000 U/mL) and PARP14i (20 nM) for 24 h. Cells were stained with an M1- ubiquitin specific antibody.

**Figure S3:**
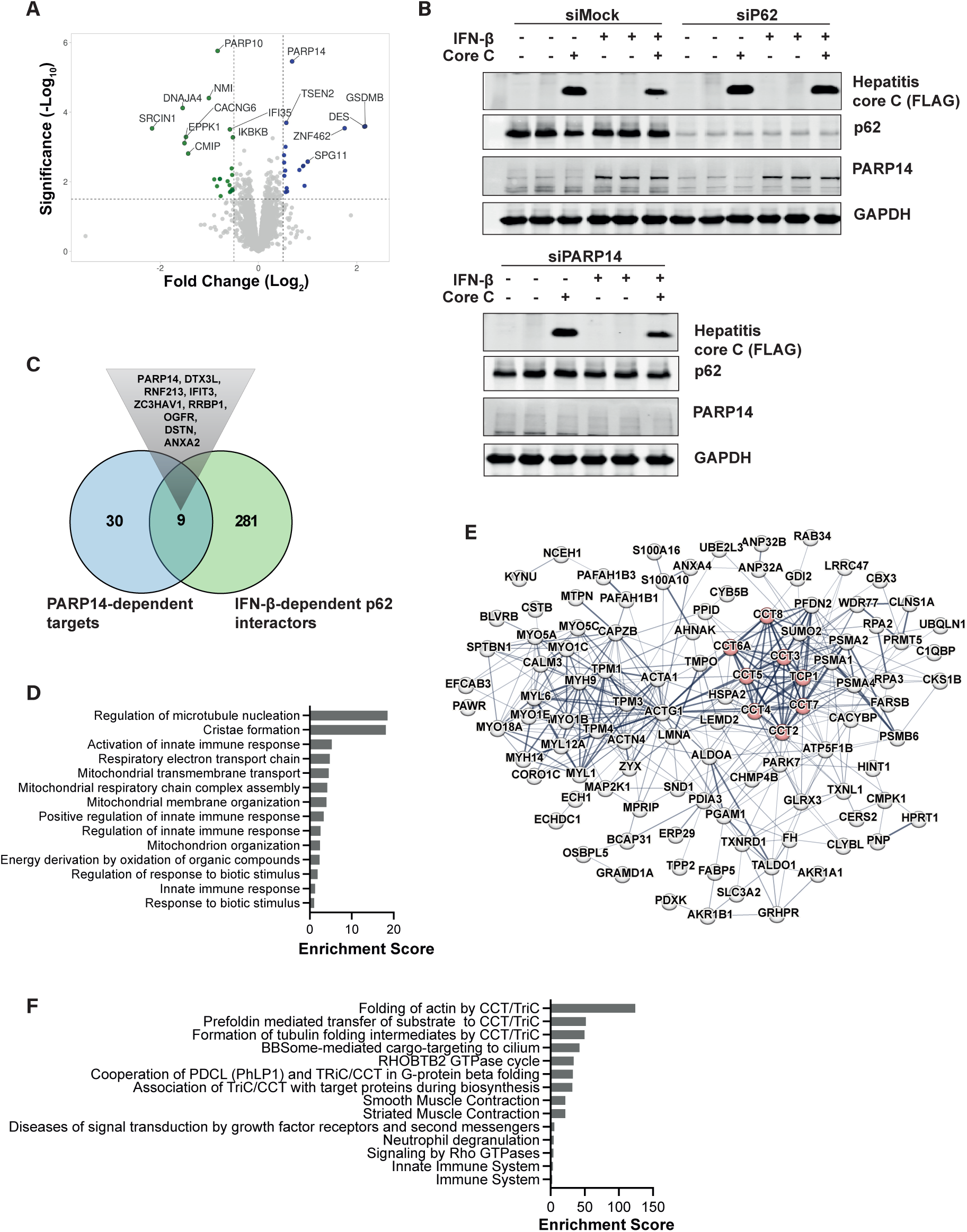
Autophagic turnover of ADP-ribosylated p62 is prevented by TRIM21. **(A)** Volcano plot showing up- and downregulated proteins after IFN-β+PARP14i treatment in comparison to IFN-β alone. Dashed lines indicate fold-change (≥1.5x change). Significance cutoff p- value (< 0.05). **(B)** Immunoblot of transiently overexpressed Hepatitis C core protein in A549 cells for 24 h. p62 and PARP14 were depleted using siRNA for 48h, and cells were treated with IFN-β (1000 U/mL) for the last 24 h of the experiment. **(C)** Overlap of the PARP14-dependent ADP-ribosylome and significantly enriched p62 interactors from our p62 co-IP MS experiment (IFN-β treated A549 cells). Proteins from the overlap are highlighted. **(D)** Bar chart showing the enrichment score (FDR < 0.05) of the GO analysis (Biological process) based on proteins from the p62 co-IP MS experiment, with proteins more enriched following IFN-β+PARP14i treatment in comparison to IFN-β alone. **(E)** STRING analysis of co-IP MS experiment showing proteins more strongly enriched following IFN-β treatment in comparison to IFN-β+PARP14i. Endogenous p62 was immunoprecipitated from A549 cells treated with IFN-β (1000 U/mL) and PARP14i (20 nM) for 24 h. Red color highlights the proteins from the TRiC/CCT chaperonin family. **(F)** Reactome pathway analysis of proteins shown in (E).

**Figure S4:**
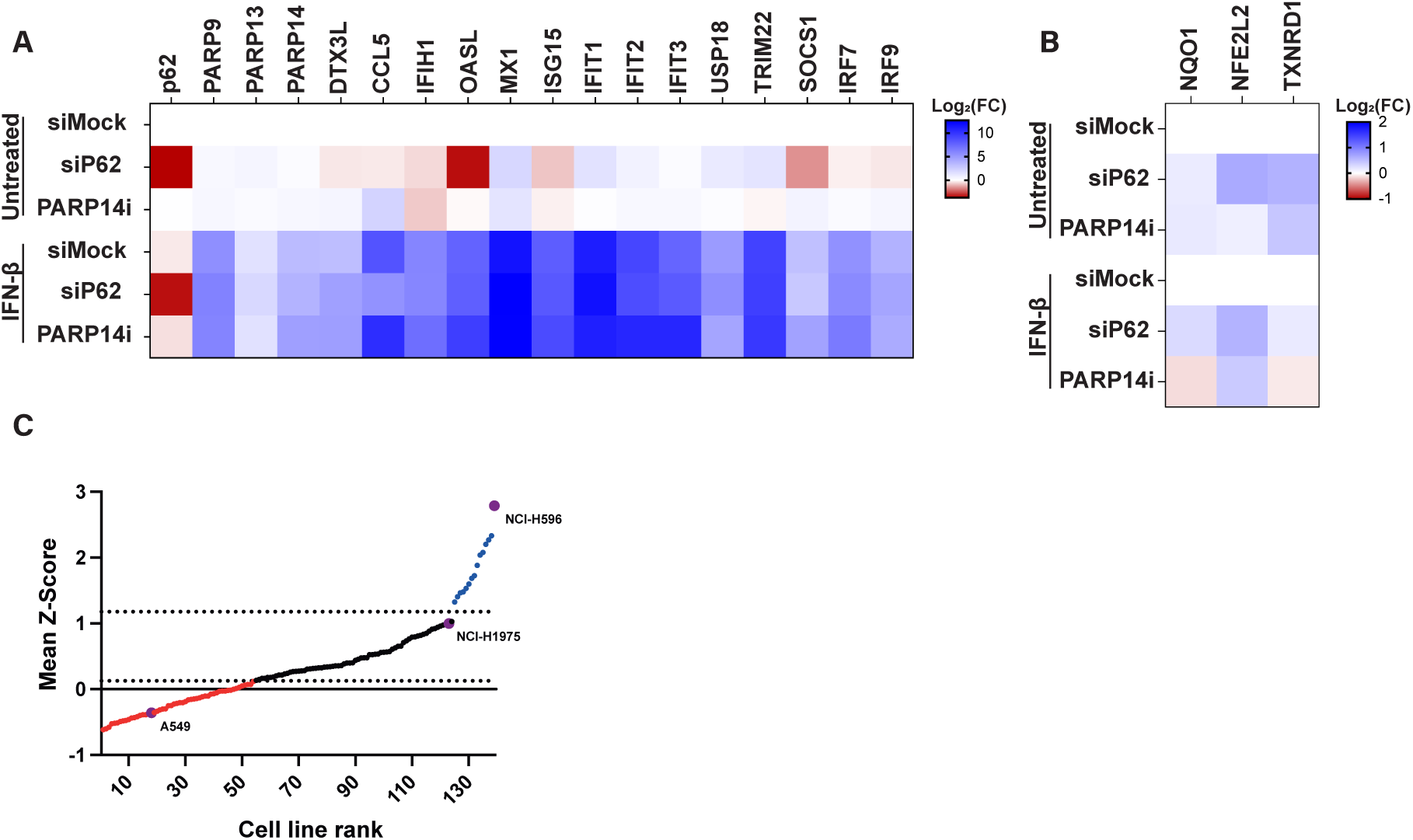
PARP14 and p62 contribute to inflammatory signaling in a context-dependent manner. **(A/B)** Heatmap showing qPCR analysis of A549 cells treated with IFN-β (1000 U/mL) and PARP14i (20 nM) for 24 h. **(C)** Scatter plot showing non-small cell lung cancer cell lines ranked by their calculated ISG score, expressed as the mean Z-score of a 38-gene ISG signature. Representative cell lines with low, intermediate, and high ISG scores are indicated, including A549, NCI-H1975, and NCI- H596, respectively.

